# Characterization of genes and alleles involved in the control of flowering time in grapevine

**DOI:** 10.1101/584268

**Authors:** Nadia Kamal, Iris Ochßner, Anna Schwandner, Prisca Viehöver, Ludger Hausmann, Reinhard Töpfer, Bernd Weisshaar, Daniela Holtgräwe

**Affiliations:** Bielefeld University, Faculty of Biology & Center for Biotechnology, Bielefeld, Germany; Julius Kühn-Institute (JKI), Institute for Grapevine Breeding Geilweilerhof, Siebeldingen, Germany

## Abstract

Grapevine (*Vitis vinifera*) is one of the most important perennial crop plants in worldwide. Understanding of developmental processes like flowering, which impact quality and quantity of yield in this species is therefore of high interest. This gets even more important when considering some of the expected consequences of climate change. Earlier bud burst and flowering, for example, may result in yield loss due to spring frost. Berry ripening under higher temperatures will impact wine quality. Knowledge of interactions between a genotype or allele combination and the environment can be used for the breeding of genotypes that are better adapted to new climatic conditions. To this end, we have generated a list of more than 500 candidate genes that may play a role in the timing of flowering. The grapevine genome was exploited for flowering time control gene homologs on the basis of functional data from model organisms like *A. thaliana*. In a previous study, a mapping population derived from early flowering GF.GA-47-42 and late flowering ‘Villard Blanc’ was analyzed for flowering time QTLs. In a second step we have now established a workflow combining amplicon sequencing and bioinformatics to follow alleles of selected candidate genes in the F_1_ individuals and the parental genotypes. Allele combinations of these genes in individuals of the mapping population were correlated with early or late flowering phenotypes. Specific allele combinations of flowering time candidate genes within and outside of the QTL regions for flowering time on chromosome 1, 4, 14, 17, and 18 were found to be associated with an early flowering phenotype. In addition, expression of many of the flowering candidate genes was analyzed over consecutive stages of bud and inflorescence development indicating functional roles of these genes in the flowering control network.

## Introduction

The reproductive developmental cycle of grapevine spans two years (S1 Figure). Grapevine plants need intense light and high temperatures to initiate inflorescences during spring, which develop and flower during the subsequent summer [1]. The ongoing tendency to higher temperatures in spring due to global warming causes earlier bud burst and flowering [2]. As a consequence, late spring frost is an increasing risk to viticulture, which may cause significant crop loss [3]. Together with flowering the onset of ripening is shifted towards earlier dates [4,5] and the ripening process occurs under warmer conditions. This influences berry composition [6], affects wine quality and promotes e.g. fungi infection. Grapevine breeding programs aim to keep the production of high quality grapes in a changing environment consistent. Making use of late flowering genotypes may be one approach to compensate for earlier ripening. Understanding the flowering process in grapevine and determining factors that lead to early or late flowering may help to control variation in berry production [7].

Detailed knowledge of pathways controlling flowering is available in crop species and the woody plant poplar, but especially the model species *A. thaliana* and rice [8,9]. With the availability of a *Vitis* reference genome sequence [10-14], gene homologs to *A. thaliana* floral development pathway genes or genes involved in photoperiod or vernalization responses could be identified in the grapevine genome. Most of these are flowering signal integrators, floral meristem identity genes, and flower organ identity genes, such as MADS box genes, like *VvMADS8* that promotes early flowering and the *VvFT/TFL1* gene family [15-17]. The expression of *VvFT*-the ortholog of the *A. thaliana FLOWERING LOCUS T* - is associated with seasonal flowering induction in latent buds and the development of inflorescences, flowers, and fruits [18]. The expression of the *LEAFY* ortholog *VvFL* is correlated with inflorescence and flower development [15]. *VvFUL-L* and *VvAP1* - homologs of the *A. thaliana* genes *FUL* and *AP1* - are suggested to act on the specification of flower organ identity as their expression appears in early developmental stages of lateral meristems and is maintained in both inflorescence and tendril primordia [16,19].

Due to the high heterozygosity and severe inbreeding depression, the first filial generation (F_1_) is used for QTL (quantitative trait loci) mapping in *V. vinifera*. This is different to other crop or model species (and is called a double pseudo test cross approach; [20,21]). Several QTL for the timing of developmental stages such as flowering time have been identified [2,22,23]. One locus contributing to flowering time control (FTC) was reported in 2006 [24]. Six QTL on different chromosomes (chr) in the mapping population GF.GA-47-42 x ‘Villard Blanc’ were described in [23]. The detected QTL are localized on chr 1, 4, 8, 14, 17, 18 and 19. Three of them (chr 1, 14 and 17) were also found in another mapping population derived from the genotypes V3125 and ‘Börner’ [23]. MADS-box genes with a proposed impact on flowering time such as *VvFL, VvFUL-L* and *VvAP1* were annotated within FTC QTL regions in *Vitis*. Further, examples of flowering time gene homologues in such QTL regions include *CONSTANS-like* genes on chr 1, 4 and 14 and the MADS-box genes, *VvFLC1* und *VvFLC2 (Vitis vinifera FLOWERING LOCUS C 1 & 2)*, which are highly expressed in buds [25].

The observation that either very early or very late flowering seems to be inherited by specific combinations of alleles at several loci, while all mixed combinations lead to an intermediate flowering type indicates an additive effect. The data further suggest a dominant effect for early flowering, with the responsible alleles being inherited from either ‘Bacchus’ or ‘Seyval’, the parents of the breeding line GF.GA-47-42 [23]. In order to link certain alleles of the sequenced genes to the flowering time phenotype, the two allele sequences of a given gene in a heterozygous diploid plant have to be determined (allele phasing).

Short read sequencing technologies still suffer from producing ambiguous haplotype phase sequences. Determining the haplotype phase of an individual is computationally challenging and experimentally expensive; but haplotype phase information is crucial in various analyses, such as genetic association studies, the reconstruction of phylogenies and pedigrees, genomic imputation, linkage disequilibrium, and SNP tagging [26,27][28,29]. In diploid organisms like grapevine, generally both alleles of a given gene are expressed. Different alleles can show different expression patterns, which can consequently result in varying manifestations of traits. The determination of these alleles is an important step in the dissection of corresponding traits. Among other approaches, haplotypic information can be obtained from DNA sequence fragments to reconstruct the two haplotypes of a diploid individual. A sequence fragment that covers at least two variant sites in a genome can link those variants together and thus phase them. When fragments are long enough to encompass multiple variant sites and the sequencing coverage is sufficiently high to provide overlaps between fragments, fragments can be assembled to reconstruct longer haplotypes [30].

For haplotype or allele phasing a variant discovery process is necessary beforehand. The two mainly used methods are based on Shotgun Genome Assembly (SGA) or on amplicon sequencing. SGA generates phasing information without knowledge of the surrounding sequence, the library coverage needs to be high and it is computationally very challenging to distinguish paralogous repeats from polymorphism but it does not require sequence information for the loci. Amplicon sequencing, which includes the amplification of a genomic region by PCR, requires sequence information of the target locus for primer design and can be done very effectively. However, it is not practical for large-scale projects [31].

In this work, we used a F_1_ population of *V. vinifera*, with the aim to associate allele sequences of several FTC candidate genes with the phenotype of flowering time in order to identify alleles influencing and controlling this trait using amplicon sequencing. Gene expression was analyzed in different time courses of bud and flower development in order to further investigate and confirm the role of FTC candidate genes.

## Materials and Methods

### Plant material

The mapping population GF.GA-47-42 x ‘Villard Blanc’ was crossed in 1989 using the breeding line GF.GA-47-42 (‘Calardis Musque’; ‘Bacchus Weiss’ x ‘Seyval’) and the cultivar ‘Villard Blanc’ (Seibel 6468 x ‘Subereux’). The 151 F_1_ individuals were planted in the vineyards at the Institute for Grapevine Breeding Geilweilerhof in Siebeldingen (49°13’05.0”N 8°02’45.0”E) in Southwestern Germany (www.julius-kuehn.de/en/grapevine-breeding) in 1996. The offspring shows notable segregation for the trait “flowering time” as the maternal breeding line GF.GA-47-42 and its parents are early flowering while the paternal line ‘Villard Blanc’ as well as its parents flower rather late. QTL analysis for flowering time was carried out using a SSR marker-based genetic map of the biparental population [32].

Phenotyping of the mapping population GF.GA-47-42 x ‘Villard Blanc’ was performed for flowering time (full bloom) in nine years (1999, 2009-2016) as described in [23] (Table 1, S1 Table). For determination of the median of flowering time for each individual, the days of the flowering period of each year were numbered whereas the first day of the flowering period was numbered with one, the second day with two, etc. These numbers were then divided by the length of the flowering period. The resulting values were used to calculate the median. Values for global radiation and accumulated temperature from November 1^st^ of the previous year until the day of full bloom were obtained from the DLR (www.wetter.rlp.de) and refer to the location of the vineyard at Siebeldingen, Germany. For gene expression analysis of FTC target genes, leaves, buds, and inflorescences from early flowering GF.GA-47-42 were collected at several consecutive time points starting from latent winter buds until inflorescences shortly before full bloom within the developmental cycle that was completed over the two consecutive years 2012 and 2013. Moreover, in 2013, sampling of buds on consecutive time points before dormancy in winter was continued. The development of the plants was described using BBCH codes [33]. Plant tissue from four different GF.GA-47-42 plants was harvested into liquid nitrogen. We decided in favor of single samples but many time points to detect trends in expression levels. Table 2 shows an overview of the collected samples.

**Table 1:** Dates of flowering periods of the mapping population GF.GA-47-42 x ‘Villard Blanc’ and the amount of global radiation at the location of the vineyards (Geilweilerhof) if available.

| Year | Start of<br>flowering<br>period<br>(days after<br>January 1<sup>st</sup> ) | End of<br>flowering<br>period<br>(days after<br>January 1<sup>st</sup> ) | Length of<br>flowering<br>period<br>(days) | Global<br>radiation at<br>beginning of<br>flowering<br>period<br>(KWh/ m<sup>2</sup>) | Global<br>radiation at end<br>of flowering<br>period<br>(KWh/ m<sup>2</sup>) |
| --- | --- | --- | --- | --- | --- |
| 1999 | 165 | 178 | 14 | / | / |
| 2009 | 156 | 170 | 15 | / | / |
| 2010 | 151 | 180 | 19 | / | / |
| 2011 | 147 | 157 | 11 | 531 | 579 |
| 2012 | 153 | 169 | 17 | 511 | 596 |
| 2013 | 168 | 183 | 16 | 516 | 597 |
| 2014 | 150 | 161 | 12 | 518 | 567 |
| 2015 | 156 | 167 | 12 | 536 | 595 |
| 2016 | 168 | 177 | 10 | 502 | 548 |

**Table 2:** Samples collected from grapevine genotype GF.GA-47-42 for the analysis of trends in gene expression levels. Listed are developmental stage and the corresponding BBCH code.

| Date of sample collection | Developmental stage | BBCH code |
| --- | --- | --- |
| Developmental cycle 2012/2013: |  |  |
| December 20<sup>th</sup>, 2012 | dormant bud | BBCH 0 |
| March 8<sup>th</sup>, 2013 | dormant bud | BBCH 0 |
| March 22<sup>nd</sup>, 2013 | swelling bud | BBCH 0-5 |
| April 12<sup>th</sup>, 2013 | swelling bud | BBCH 5-9 |
| April 26<sup>th</sup>, 2013 | swelling bud/ first leaves | BBCH 11 |
| May 3<sup>rd</sup>, 2013 | buds/ first leaves | BBCH 11-13 |
| June 7<sup>th</sup>, 2013 | inflorescences & leaves | BBCH 53 |
| June 14<sup>th</sup>, 2013 | inflorescences & leaves | BBCH 55 |
| June 17<sup>th</sup>, 2013 | inflorescences | BBCH 57 |
| Developmental cycle 2013/2014: |  |  |
| July 22<sup>nd</sup>, 2013 | buds & leaves | / |
| August 2<sup>nd</sup>, 2013 | buds | / |
| August 8<sup>th</sup>, 2013 | buds & leaves | / |
| August 16<sup>th</sup>, 2013 | buds | / |
| August 22<sup>nd</sup>, 2013 | buds & leaves | / |
| September 5<sup>th</sup>, 2013 | buds | / |
| September 19<sup>th</sup>, 2013 | leaves | / |

### FTC candidate gene prediction

For the identification and characterization of putative flowering time control (FTC) genes, functional data from well studied model species was used to exploit the grapevine genome for homologous genes. Using BLAST (e-value cut off below 1e-25) [34] protein sequences of candidate genes from *A. thaliana* and other model species were compared against the *Vitis* protein sequences (PN40024-12xv0, Genoscope gene prediction 12X.v0 (www.genoscope.cns.fr/externe/GenomeBrowser/Vitis/) and the CRIBI gene prediction 12X.v2 [12]). Results were manually checked for additional evidence from the literature.

For functional annotation of FTC candidate genes, the method of reciprocal best hits (RBH) [35] was applied. A RBH pair consists of two sequences from different sets of sequences, each displaying the highest genome wide score in the other data set. Genomic sequences of FTC genes were compared against protein sequences of *V. vinifera* and *A. thaliana* with blastx. If a gene displayed several transcripts, the longest sequence was used. Using tblastn the hit showing the highest score was compared back against *V. vinifera* coding genes. When the original query was found to have the highest score, the resulting RBH pair was considered.

To establish unique genes, we used the *Vv* (*Vitis vinifera*) prefix followed, for almost all genes, by the gene name deduced from the *Arabidopsis* annotation. In many cases the *Vitis* genome holds several putative homologs for known FTC genes from model crops, leading to low number of RBHs between *Vitis* and *Arabidopsis* genes. In order to distinguish these *Vitis* genes, the one with the highest BLAST score to the query gene got the name extension “a”, the second best the “b”.

### Amplimer design

Genes for targeted allele phasing (target genes) through amplicon sequencing were selected out of the identified FTC candidate genes. The cDNA sequences of target genes were used as query in a BLAST against the grapevine reference sequence PN40024-12xv0. Genomic DNA sequences were extracted in addition to 1,000 bp from the 5’-and 3’-UTR regions. Primers were designed for overlapping amplimers of up to 8 kb using the tool Primer3 [36].

### DNA isolation and amplicon generation

Extraction of genomic DNA was performed from young leaf tissue. The leaf material was grounded under liquid nitrogen and subsequently used for DNA isolation with the DNeasy® Plant Maxi Kit (Qiagen, Hilden, Germany) according to manufacturer’s protocols. The purified DNA was quality checked via gel electrophoresis and quantified using a NanoDrop spectrophotometer (Peqlab, Erlangen, Germany). Amplicons were amplified by long range PCR (98 °C 30 sec, 15 cycles of 10 sec 98 °C, 30 sec 72 °C – 57 °C, 5 min 72 °C, 25 cycles 10 sec 98 °C, 30 sec 58 °C, 5 min 72 °C and finally 2 min 72 °C).

Target gene sequences were amplified from 37 individuals of the mapping population GF.GA-47-42 x ‘Villard Blanc’ including the parental lines and 35 F_1_ individuals with early, intermediate and, late flowering time phenotypes (S2 Table).

### Library preparation and amplicon sequencing

Amplicon sequencing was carried out on a MiSeq (Illumina, San Diego, USA) in seven runs. All amplicons belonging to a respective individual were pooled in equimolar amounts, fragmented by sonification using a Bioruptor (Diagenode, Denville, USA) and subsequently used for library preparation. The libraries were prepared as recommended by Illumina (TruSeq DNA Sample Preparation v2 Guide). Adaptor-ligated fragments were size selected on a two percent low melt agarose gel to an average insert size of 500 bp. Fragments that carry adaptors on both ends were enriched by PCR. Final libraries were quantified using PicoGreen (Quant-iT, Fisher Scientific, Schwerte, Germany) on a Fluostar platereader (BMG labtech, Ortenberg, Germany) and quality checked by HS-Chips on a 2100 Bioanalyzer (Agilent Technologies, Santa Clara, USA). Up to 20 libraries were pooled and sequenced on an Illumina MiSeq platform with 2 x 250 bp read length using the Illumina MiSeq v2 reagents. After sequencing, basecalling and demultiplexing and FASTQ file generation was performed using a casava-based in house script.

### Read processing and mapping

Adapter trimming of raw reads and quality filtering of reads with a window of four consecutive bases that exhibited a quality value below 30 was performed using Trimmomatic [37]. Bases at the heads and tails of the reads with quality values below 30 were cropped using Trimmomatic. Before and after trimming the tool FastQC (www.bioinformatics.babraham.ac.uk/projects/fastqc) was used to check the quality of the reads. Between 11.5 and 35.6% (20.2% on average [standard deviation (SD): 5.5%]) of reads were dropped through trimming. Trimmed reads were mapped to the grapevine reference sequence PN40024 12x.v2 [14] using the BWA-MEM algorithm which is suitable for long reads with default parameters [38]. Mapping was performed for each individual separately. Instead of the entire reference sequence the target gene sequences only were chosen for mapping in order to prevent false positive mapping results. The SAM format files were converted to BAM format files and sorted using SAMtools [39]. Readgroups were added and duplicated reads removed using Picard Tools (https://broadinstitute.github.io/picard/). Besides PCR duplicates unpaired reads were removed from the mapping files. About 15% of amplicons failed to be amplified or sequencing depth was below 20.

### Allele phasing of target genes

In order to separate the two alleles of the sequenced target genes (phasing), a workflow using the Genome Analysis Toolkit (GATK) [40] was established (Fig. 1). After read alignment, the quality of the alignments was improved in two ways. Firstly, local realignments around InDels were performed using InDelRealigner of GATK [40] to reduce the number of misalignments. Occasionally, the presence of insertions or deletions in individuals with respect to the reference genome sequence leads to misalignments of reads to the reference, especially when InDels are covered at the start or end of a read. Such misalignments lead to many false positive SNPs. Secondly, base quality scores of reads in the aligned mapping files were recalibrated using BaseRecalibrator of GATK in order to correct for variation in quality with machine cycle and sequence context. Thus, more accurate and more widely dispersed quality scores are provided.

**Fig. 1:**
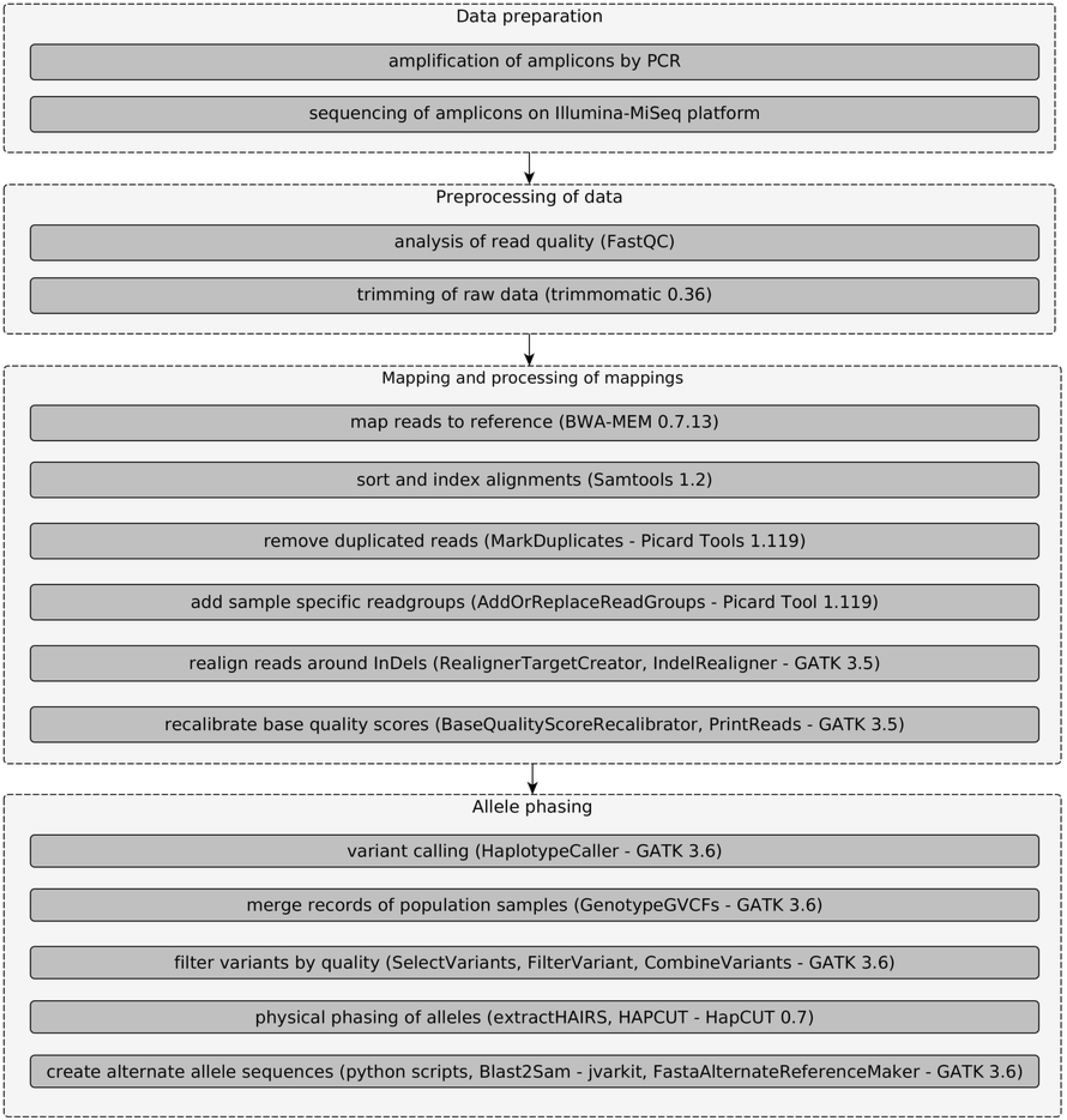
Workflow using the Genome Analysis Toolkit (GATK), which uses the high-coverage genotype sequence variation information and the family relationship for phasing.

Using the HaplotypeCaller of GATK variants were called for each individual separately. The ploidy parameter was set to 12 for variant calling. It was performed in gVCF mode for F_1_ individuals and the parental lines of the population GF.GA-47-42 x ‘Villard Blanc’. Cases of allele dropout were identified, in which the missing allele leads to genotyping errors. Since we were working with an F_1_ population and by applying Mendelian constraints it was possible to determine which allele was missing within the population GF.GA-47-42 x ‘Villard Blanc’, but its sequence remained unknown. After variant calling, resulting variant files from individuals of the population were merged using GATK’s GenotypeGVCFs in order to apply further downstream steps on all samples together. At each position of the input gVCFs, this tool combines all spanning records and outputs them to a new variant file. Raw variants were hard-filtered according to GATK’s “Best Practices” recommendations [41,42]. In addition, variants with read coverage depth and genotype quality below 20 were filtered out. For the determination of allele-specific sequences initially physical phasing was performed using HapCUT [30]. Fragments were defined from the sequenced reads. Haplotype-informative reads that cover at least two heterozygous variants were extracted from the aligned file using the tool extractHairs from HapCut and used for the assembly of haplotypes. The information of polymorphic sites was passed to HapCUT through a variant file. A maximum number of 600 iterations were used to run HapCut and the reference sequence was provided in order to extract reads covering both SNPs and InDels. Using various python scripts, intervals in which phasing could be performed in individuals of the population GF.GA-47-42 x ‘Villard Blanc’ including the parents and F_1_ individuals were determined and homozygous alternative variants were added to the variant files. Using GATKs FastaAlternateReferenceMaker FASTA-format files with alternate sequences were created for each individual within the regions in which allele phasing could be performed.

A nomenclature system was created for the alleles of genes within the population GF.GA-47-42 x ‘Villard Blanc’ (S3 Table). The system distinguishes between fourteen different cases, where four, three, or two different allele sequences can be present at a locus or all sequences can be identical. Moreover, it distinguishes between various combinations of two or three different sequences. E, as in E1, E2 and E0, refers to “early” and originates from early flowering GF.GA-47-42, while L, as in L1, L2 and L0 refers to “late” and originates from late flowering ‘Villard Blanc’. N means that both GF.GA-47-42 and ‘Villard Blanc’ share one or more alleles. N1 means that E1 and L1 are alike, while N2 means that E2 and L2 are alike. N means that either L2 and E1 or E2 and L1 are alike. Na means that E1, E2, and L1 are alike. Nb means that E1, E2, and L2 are alike. Nc means that E1, L1, and L2 are alike. Nd means that E2, L1, and L2 are alike. Descriptions for allele combinations that distinguish between which of the two alleles of one parental line is alike the two alleles of the other line (as in NaNa x NaL2) was implemented in order to be able to track patterns of allele combinations throughout QTL regions and closely neighboring genes.

### Correlation analysis

To test for the correlation of an allele and the flowering time phenotype, a Wilcoxon Rank-Sum test was carried out between a dichotomous variable (the presence or absence of an allele) and a continuous variable (flowering time). The null hypothesis assumed that the median of flowering time between groups of individuals carrying a certain allele or not is equal. When p-values below 5% were found, the null hypothesis was rejected and an association between an allele and the flowering time phenotype was found to exist.

### Marker development and testing of the whole mapping population

After creating haplotype specific allele sequences through amplicon sequencing and the subsequent bioinformatic pipeline, markers were designed for haplotype specific PCRs. Obtained allele sequences of target genes were scanned for InDel structures differing between the parental alleles. Variants with low coverage or low quality were filtered out. In the case that InDels were filtered out, the actual allele sequence can be greater than the calculated one. The sequence information was used for subsequent STS (Sequence-Tagged Sites) marker design with the Primer3 tool [36]. Primers had an optimum Tm of 58-60 °C, with PCR products differing in size between 100 – 400 bp for multiplexing purposes (S7 Table). Forward primers were labeled at the 5’end with one of the fluorescent dyes 6-FAM (blue), HEX (green), TAMRA (yellow) or ROX (red). Allele distributions were analyzed over all 151 F_1_ individuals of the mapping population GF.GA-47-42 x ‘Villard Blanc’. PCRs were carried out with the QIAGEN multiplex PCR kit (Qiagen GmbH, Hilden, Germany) following the instructions of the manufacturer in three multiplexes combining different product sizes and fluorescent dyes. Resulting PCR products were analyzed on an ABI 3110xl Genetic Analyzer (Applied Biosystems, Foster City, USA) and the results compared with the respective phenotype of the tested individual (i.e. early, intermediate or late flowering).

### RNA extraction and sequencing

Total RNA was extracted from up to 100 mg of liquid nitrogen ground tissue using the Spectrum™ Plant Total RNA kit (Sigma-Aldrich, Taufkirchen, Germany) according to the manufacturer’s instructions for protocol B. After on-column DNase treatment with the DNase I Digest Set (Sigma-Aldrich, Taufkirchen, Germany) the RNA was quantified. RNA-libraries for each time point were prepared according to the Illumina TruSeq RNA Sample Preparation v2 Kit using an input of 1 μg of total RNA. RNA-Seq (1x 135 bp) was performed on an Illumina Rapid HiSeq-1500 Run. One barcoded library was created for each of the time points.

### RNA-Seq read processing for analysis of gene expression kinetics

Read trimming and quality control was performed as described above in “Read processing and mapping”. Sequence read data are available from SRA accession SRP153932. The reads were mapped to the grapevine reference sequence PN40024 12x.v2 [14] using tophat2 [43] which is capable of performing split read mapping. The maximal intron size was set to 3000, otherwise default parameters were used. Resulting BAM-format files were sorted and indexed using SAMtools [39]. With HTSeq [44] mapped reads were counted for each gene. Differential gene expression was analyzed using the R-package DESeq2 [45]. In order to perform an analysis of expression without replicates, the counts were modeled as a smooth function of time, and an interaction term of the condition with the smooth function was included. Likelihood ratio test of DESeq2s with a reduced design, which does not include the interaction term, was then applied. Genes with small p-values from this test are those showing a time-specific effect.

## Results

### Phenotypic evaluation of the mapping population

The 151 F_1_ individuals of the segregating population and their parental lines were phenotyped for time of full bloom as indicated in S2 Figure showing the timing of flowering in days after January 1^st^. The length of the flowering period varied considerably between 10 days (2016) and 17 days (2012) [23]. In the year 2010 the flowering period was heavily extended compared to the other years. The greatest portion of individuals within the population reached full bloom in approximately the first third of the flowering period. Within the mapping population, early flowering is inherited from the maternal genotype GF.GA-47-42.

### Identification of FTC candidate genes

Functional data from *A. thaliana* and other model organisms was systematically exploited to identify FTC candidate genes in the *Vitis* reference genome sequence. More than 500 homologous genes were identified which are distributed over all chromosomes including the unanchored, random part of the sequence (S4 Table). Some of the genes are absent from the CRIBI annotations, but were included in the previous annotations, provided by Genoscope. To our knowledge the majority of the identified FTC candidate genes was not analyzed or even mentioned in a previous publication. As expected, an enrichment of the FTC candidate genes (75) annotated within the FTC QTL regions was found. In several cases we identified more than one homologous sequence in the grapevine genome with a single copy *Arabidopsis* query. In these cases not necessarily the gene with the highest sequence similarity is the one in the FTC QTL region, nor the one with the highest expression in flowering related tissues. For instance the *RAV* genes *VvRAV1b* and *VvRAV1c* are located within the QTL regions on chr 1 and chr 14, respectively, whereas the *RAV1a* is located on chr 11 outside of any FTC QTL.

Many of the FTC candidate genes are transcription factors involved in flower development and morphogenesis such as members of the AP2/EREBP family [46] and homeodomain proteins [47]. About eight MYB-transcription factors that participate in cell cycle control in many living taxa [48] were among the identified FTC candidate genes in *Vitis*. Several other protein families were among the FTC candidate genes, such as a dozen GRAS and FRIGIDA proteins that are involved in flowering time and plant development. FRIGIDA proteins are required for the regulation of flowering time by upregulating *FLC* expression. Allelic variation at the FRIGIDA locus is an important determinant of natural variation in the timing of flowering [49]. The GRAS (GAI, RGA, SCR) family is a very important family of proteins involved in flowering in grapevine. GRAS proteins participate in GA signaling, which influences numerous aspects of plant growth and development [50]. Remarkably sixteen SQUAMOSA PROMOTER BINDING PROTEIN (SBP)-domain proteins, that are known from other plants as transcriptional activators involved in a variety of processes such as flower and fruit development, plant architecture, GA signaling, and the control of early flower development [51] are candidates.

### Allele phasing

From our comprehensive list of *V. vinifera* FTC candidates the 72 most promising genes were chosen as targets for amplicon sequencing (S5 Table), many of which are located in flowering related QTL regions on chr 1, 14, and 17 [23]. The average read depth of coverage was 286 (SD: 276) and for most samples sequencing depth was between 100 and 300. Variants in the analyzed lines were detected with a density between 1.02 and 1.63 variants per 100 bp most of which were SNPs.

In order to link certain alleles of the sequenced candidate genes to the flowering time phenotype, the two alleles of genes had to be reconstructed from the mix of sequenced fragments of the two alleles. The phasing of alleles was performed on the basis of sites polymorphic between the two alleles of a gene.

Aside from recombination events, a parent-offspring pair must share one haplotype for each chromosome and thus one identical-by-descent allele for every gene. Hence, Mendelian constraints could be applied to validate the obtained allele-specific sequence. Alleles of the chosen 72 target genes studied could be identified in 46 cases (S5 Table).

In 23 cases four different allele sequences could be found, three allele sequences in 18 cases, two in four cases and in one case (VIT_217s0000g00150; *VvFL*) only one allele sequence, meaning that all individuals of the population were homozygous for the respective locus. This fits the expectation since grapevine is highly heterozygous. The number of allele sequences has been deduced from regions of the genes in which phasing was performed. The lengths of the phased intervals were between 204 and 8,285 bp (S5 Table).

### Correlation analysis

Allele sequences of the progeny of the mapping population GF.GA-47-42 x ‘Villard Blanc’ were compared against the allele sequences of the parental lines to determine the inheritance pattern within the population for each gene. In order to find alleles correlating with the phenotype of flowering time, a correlation analysis between the phased alleles of FTC target genes and flowering time phenotypes was performed. Several sets of phenotypic data were used. For the years 1999, 2009-2016 a correlation analysis was performed using days after January 1^st^ of the respective year. Additionally for the years 2011-2016 values of accumulated temperature above 3°C from November 1^st^ of the previous year and global radiation in KWh/m^2^ from January 1^st^ were considered.

After the reconstruction of inheritance patterns within the parental lines and the 35 analyzed F_1_ individuals of the mapping population GF.GA-47-42 x ‘Villard Blanc’ through the amplicon sequencing approach and subsequent bioinformatic analysis, the numbers of individuals harboring each of the alleles was determined and a correlation analysis between alleles of FTC target genes and the flowering time phenotype was performed for 43 genes. A correlation between alleles and flowering time could be observed for several genes on chr 1, 4, 14, 17, 18, and within unassigned contigs. Correlation values differed depending on whether days, accumulated temperature or global radiation was used as phenotypic data. As an example Fig. 2 shows allele combinations in the parental lines of the population GF.GA-47-42 x ‘Villard Blanc’ and the p-values of the correlation of alleles unique to one of the lines. Values equal and below 0.05 were considered to be significant and the lower the p-value the higher is the correlation. In total for 16 FTC target gene alleles a significant correlation with either an early or late flowering phenotype could be found.

**Fig 2:**
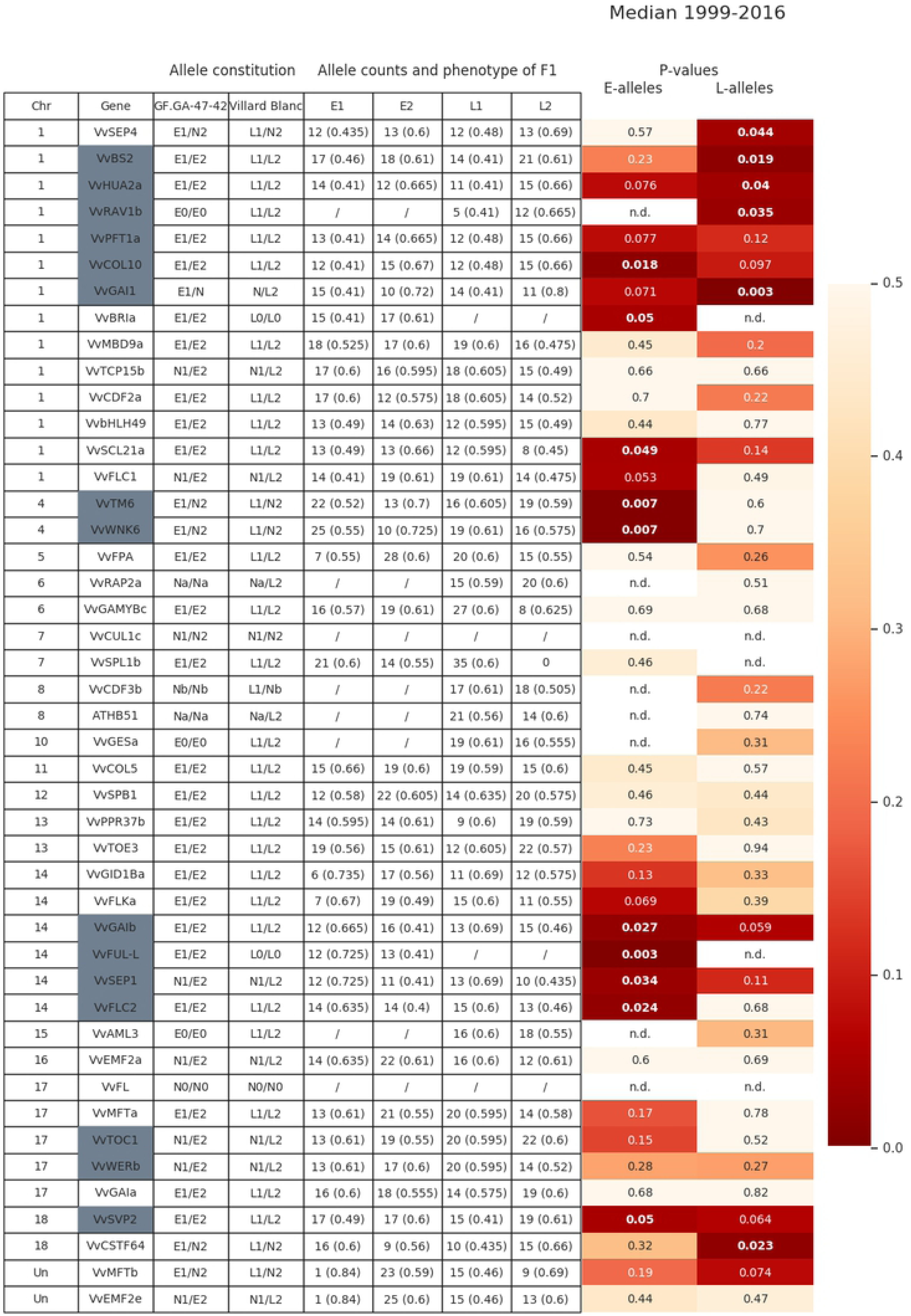
Correlation between alleles of FTC target genes and flowering time phenotype. Given are the allele constitutions of the parental lines for each gene and the allele counts of the amplicon sequenced F_1_ individuals. The median of flowering time (calculated from days after January 1^st^ of the years 1999 and 2010-2016) of individuals carrying the counted is given in brackets. The higher the value of the median, the later the flowering phenotype of the F_1_ individuals. Color coded are the p-values for the E alleles and L alleles in the up to 35 F_1_ individuals. Significant correlation values are in bold and italic. Genes located in QTL regions are marked in grey. Differences in allele counts between the years are due to missing data points. “E” alleles are inherited from GF.GA-47-42, while “L” alleles originate from ‘Villard Blanc’. “N” means that both GF.GA-47-42 and ‘Villard Blanc’ share one or more alleles. “E0”: E1=E2, “L0”: L1=L2, “N1”: E1=L1, “N2”: E2=L2. “N”: L2=E1 or E2=L1, “Na”: E1=E2=L1, “Nb”: E1=E2=L2. "n.d.”: not determined. Further explanations are given in S3 Table.

The L2 alleles, inherited from the paternal line ‘Villard Blanc’, of *VvSEP4 (SEPALLATA 4), VvBS2, VvHUA2a, VvRAV1b*, and *VvGAI1* (chr 1) correlate with late flowering, strengthen the importance of the FTC QTL on chr1. The E1 alleles of the two genes *VvWNK6 (V. vinifera WITH NO LYSIN KINASE 6)* and *VvTM6* (*V. vinifera TOMATO MADS-BOX 6*), both located on chr 4 and inherited from the early flowering maternal line, were found to strongly correlate with early flowering. The p-values calculated from the median (Fig 2 is p = 0.007 and values down to p = 0.003 were observed for single years. Table 3 shows the p-values of correlation for different sets of phenotypic data related to *VvWNK6* and *VvTM6*. Most of the significant correlations are obvious regardless the year or scale of phenotyping (days after January 1^st^, accumulated temperature or global radiation). The differences in correlation among years are due to the seasonal weather conditions of the respective year, which influence both the flowering time and the length of the flowering period. A significant correlation between the E1 allele of Vv*WNK6* and the flowering time phenotype could not be observed in 2016 for neither days after January 1^st^, accumulated temperature or global radiation. In 2015, the correlation was not significant for days after January 1^st^ but, albeit only slightly, for the other two sets of phenotypic data. Other genes, such as *VvMFT* (*V. vinifera MOTHER of FT and TFL1*) showed significant correlation in 2016 but not in 2013.

**Table 3:** P-values of the correlation between the E1 allele distribution of *VvWNK6* and *VvTM6* in relation to different sets of phenotypic data using 35 amplicon sequenced F_1_ individuals.

|  | ***VvWnk6* (E1)** | ***VvTM6* (E1)** |
| --- | --- | --- |
| Days after January 1<sup>st</sup>/ 1999 | 0.032 | 0.023 |
| Days after January 1<sup>st</sup>/ 2009 | 0.012 | 0.009 |
| Days after January 1<sup>st</sup>/ 2010 | 0.44 | 0.37 |
| Days after January 1<sup>st</sup>/ 2011 | 0.033 | 0.063 |
| Days after January 1<sup>st</sup>/ 2012 | 0.047 | 0.041 |
| Days after January 1<sup>st</sup>/ 2013 | 0.008 | 0.012 |
| Days after January 1<sup>st</sup>/ 2014 | 0.015 | 0.029 |
| Days after January 1<sup>st</sup>/ 2015 | 0.067 | 0.063 |
| Days after January 1<sup>st</sup>/ 2016 | 0.177 | 0.098 |
| Median for days after January 1<sup>st</sup>/ 1999-2016 | 0.012 | 0.009 |
| Accumulated Temp. above 3°C/ 2011 | 0.027 | 0.109 |
| Accumulated Temp. above 3°C/ 2012 | 0.03 | 0.091 |
| Accumulated Temp. above 3°C/ 2013 | 0.004 | 0.058 |
| Accumulated Temp. above 3°C/ 2014 | 0.003 | 0.016 |
| Accumulated Temp. above 3°C/ 2015 | 0.046 | 0.186 |
| Accumulated Temp. above 3°C/ 2016 | 0.177 | 0.098 |
| Global radiation (KWh/m<sup>2</sup>)/ 2011 | 0.027 | 0.109 |
| Global radiation (KWh/m<sup>2</sup>)/ 2012 | 0.03 | 0.091 |
| Global radiation (KWh/m<sup>2</sup>)/ 2013 | 0.004 | 0.058 |
| Global radiation (KWh/m<sup>2</sup>)/ 2014 | 0.003 | 0.016 |
| Global radiation (KWh/m<sup>2</sup>)/ 2015 | 0.046 | 0.186 |
| Global radiation (KWh/m<sup>2</sup>)/ 2016 | 0.177 | 0.098 |

Compared to the reference sequence, the E1 allele of *VvWNK6* (chr 4) was found to harbor a variation in the terminal exon (SNP at chr4:21997435/ C → T) leading to an amino acid exchange from threonine to methionine. Fig. 3 shows the distribution of allele combinations for *VvWNK6* among individuals of the mapping population. Early flowering is associated with the E1 allele inherited from the maternal ‘Bacchus’ allele of GF.GA-47-42.

**Fig 3:**
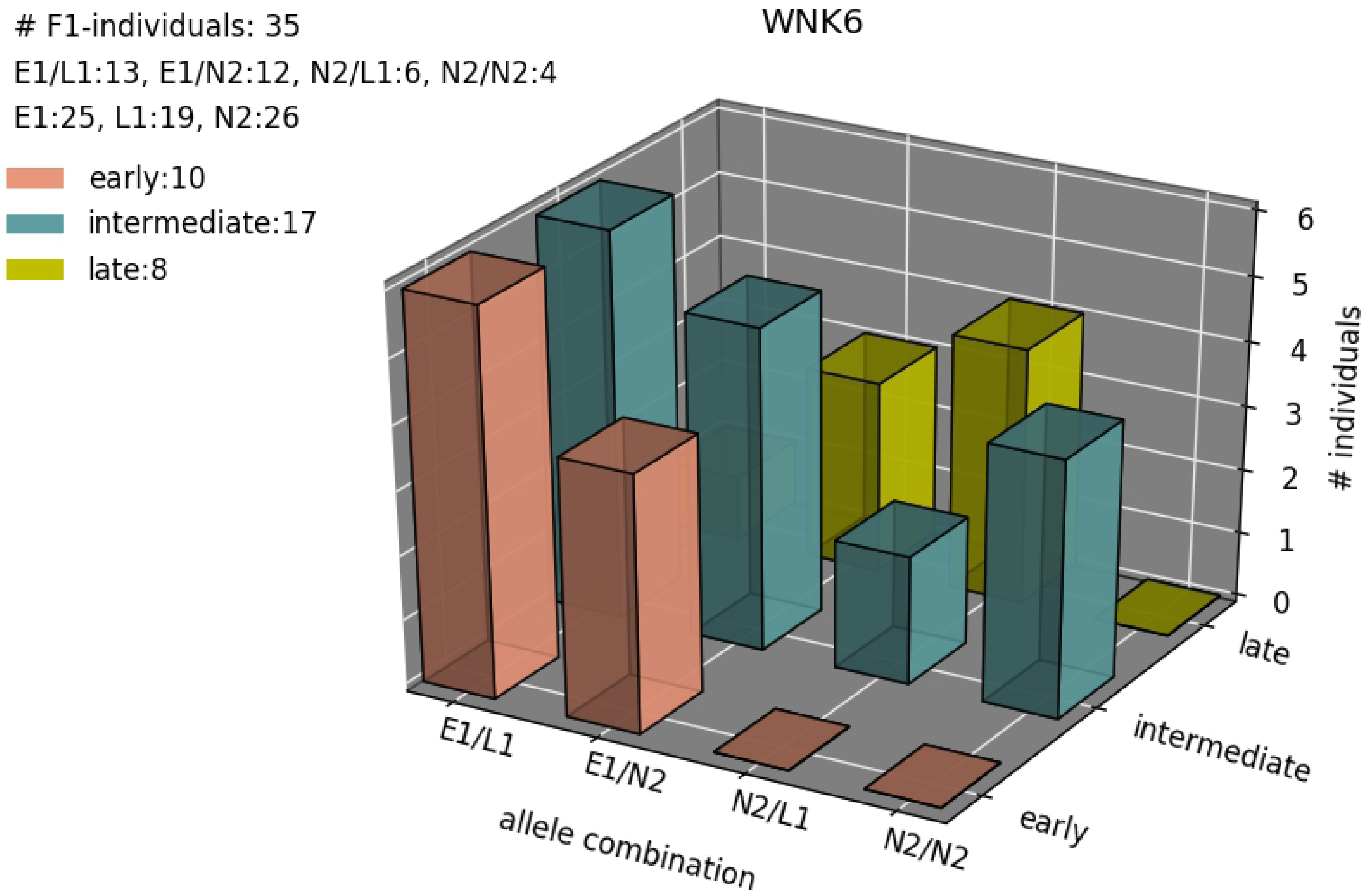
Distribution of allele combinations for *VvWNK6* (chr 4) among 35 selected individuals of the mapping population GF.GA-47-42 x ‘Villard Blanc’. The date of flowering was counted in days from the 1^st^ of January and the data was subsequently classified according to six stages for flowering time following (1 = very early flowering; 2 = early flowering; 3 = medium early flowering; 4 = medium late flowering; 5 = late flowering; 6 = very late flowering). For visualization flowering classes 1 and 2, 3 and 4, and 5 and 6 were merged.

### Application of the pipeline for amplicon sequencing in a heterozygous plant for subsequent marker design

Amplicon sequencing was performed in 35 F_1_ individuals and the parents of the mapping population. In order to investigate the resulting allele distributions over all 151 F_1_ individuals of the mapping population GF.GA-47-42 x ‘Villard Blanc’, STS markers were designed from the allele sequences that enabled an easy allele-specific genotyping. The information obtained from amplicon sequencing of the FTC target genes proved usable for both deduction of segregation patterns and marker design for investigating allele distribution over the whole mapping population. Table 4 gives an overview of the segregation patterns as analyzed for all 151 F_1_ individuals. From 15 markers 12 showed a segregation pattern matching the segregation pattern that was obtained through allele phasing. The markers GAVBInd_019 and GAVBInd_020 were not designed using the obtained allele sequences of GF.GA-47-42 and ‘Villard Blanc’, since suitable InDels were not available. Therefore, these markers were designed based on InDels upstream of the phased regions. Observed product sizes can deviate from the expected ones by 1 – 2 bp due to the limited accuracy of the used fragment analyzing method. Markers GAVBInd_004, GAVBInd_014, and GAVBInd_019 showed two different segregation patterns since the measuring method cannot reliably resolve differences of 1 – 2 bp. See S6 Table for further details.

**Table 4:**
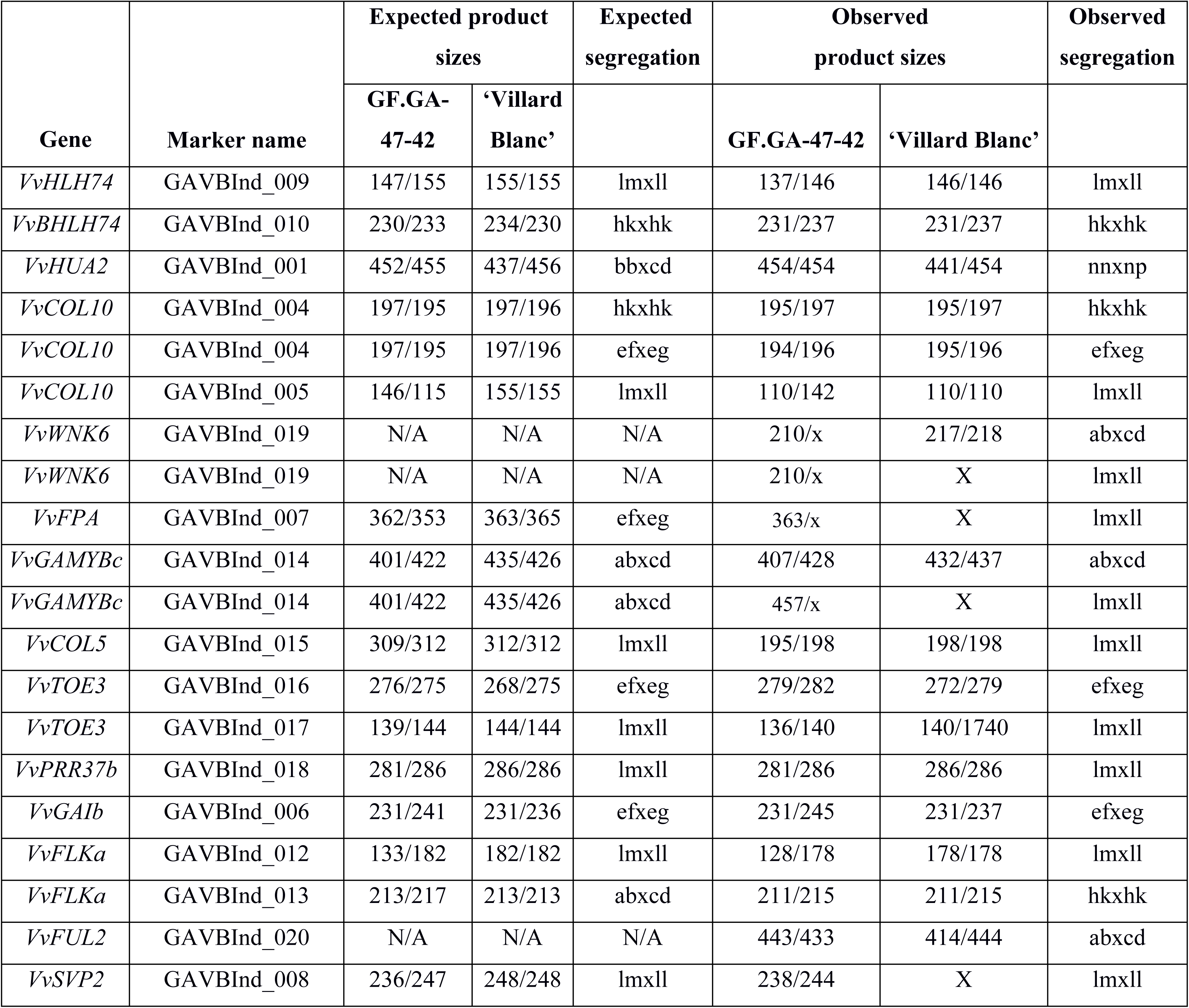
Comparison of the expected and observed allele sizes (bp) and segregation patterns of several FTC target genes. Expected data were obtained through amplicon sequencing; observed data were gained by analyzing 151 F_1_ individuals of the mapping population GF.GA-47-42 x ‘Villard Blanc’ with STS markers located within the FTC target genes. ab x cd: four alleles/both parents heterozygous, hk x hk: 2 alleles/both parents heterozygous, ef x eg: 3 alleles/both parents heterozygous, lm x ll: 2 alleles/ mother heterozygous, nn x np: 2 alleles, father heterozygous. x: amplification failed. See Table S6 for further information.

Using the results of marker segregation across the 151 F_1_ individuals, a correlation analysis between alleles and flowering time phenotypes was performed. The correlation results of marker analysis support those of allele phasing (Table 5). See S7 Table for further details.

**Table 5:**
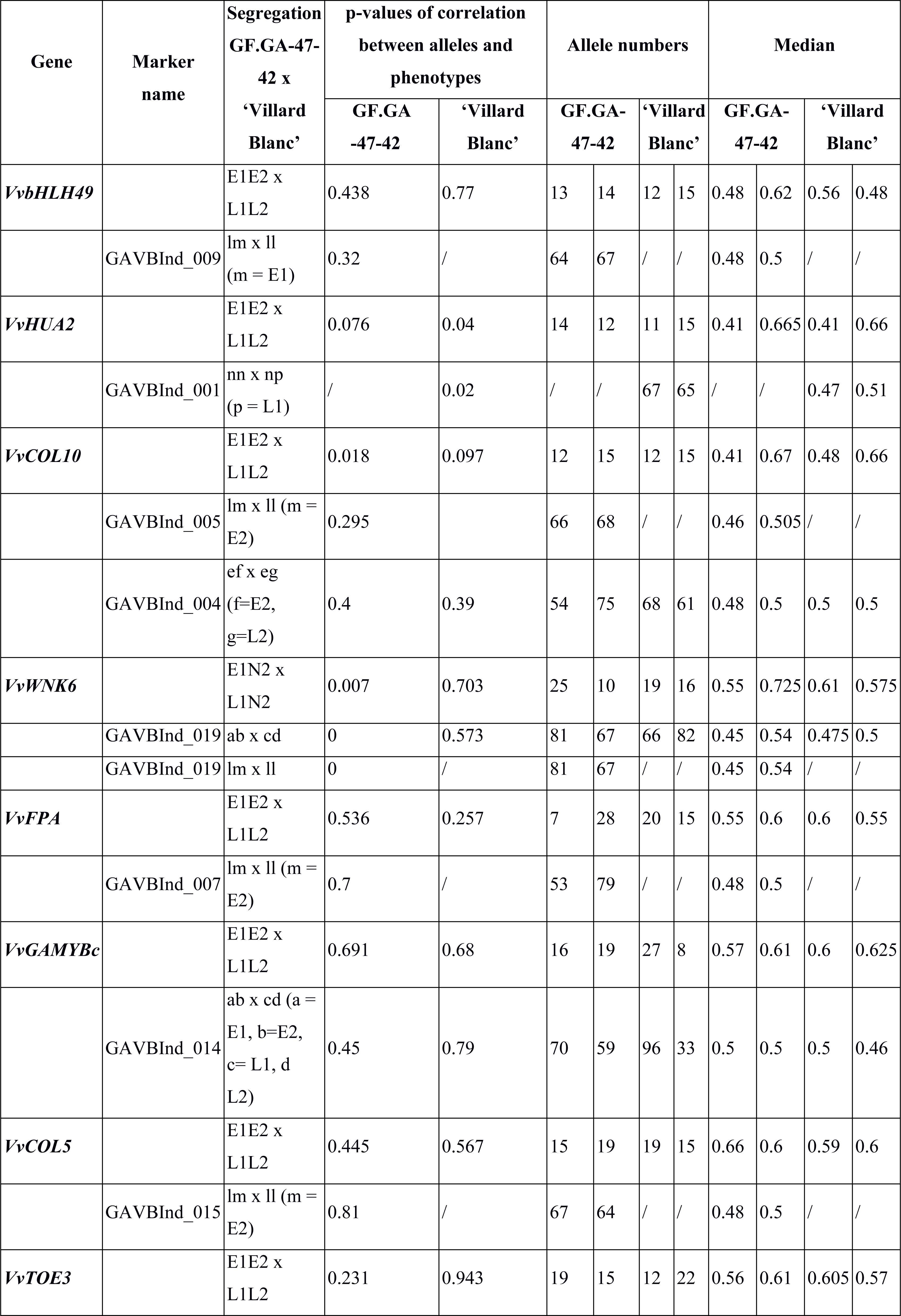
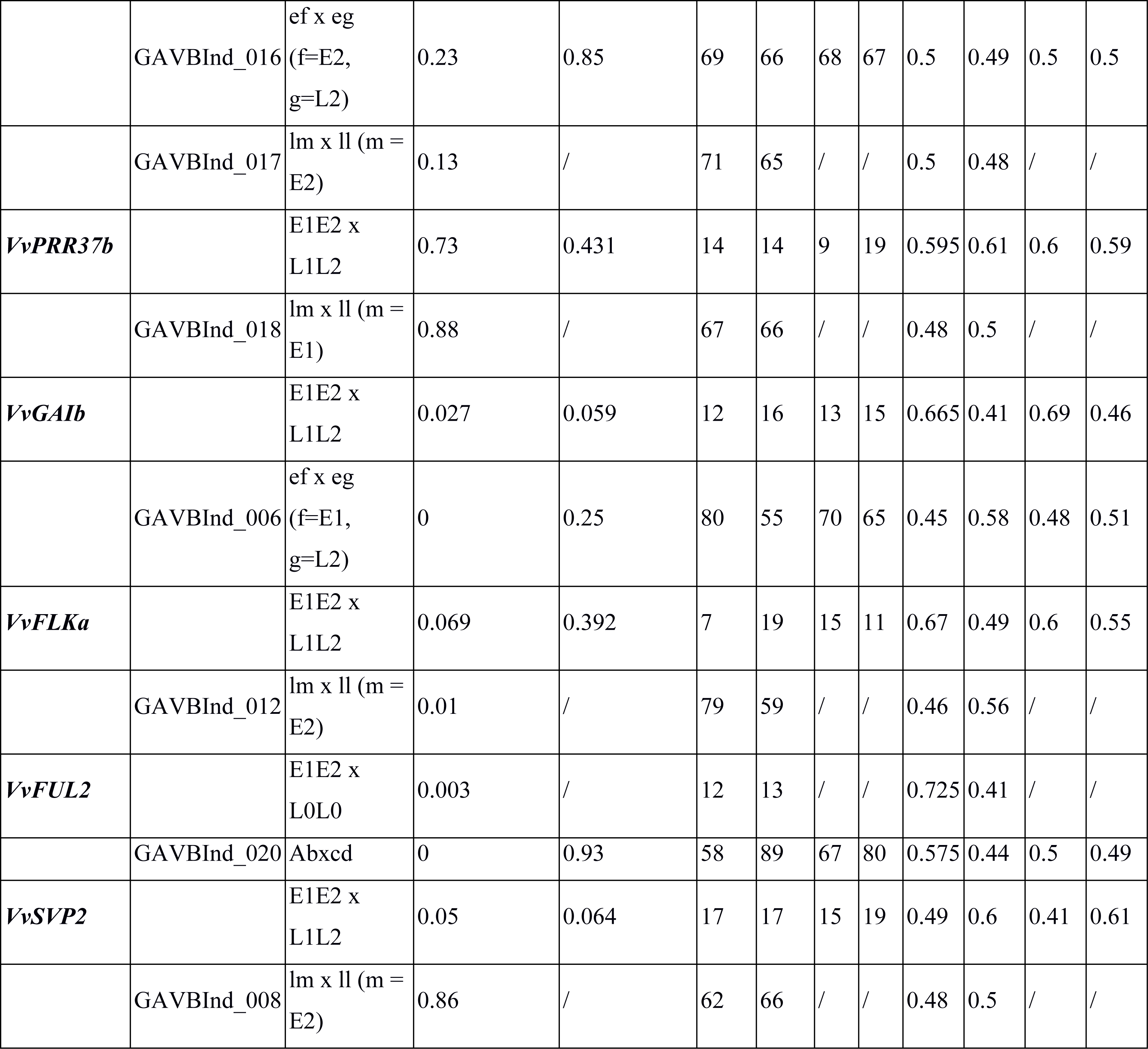
P-values of correlation between alleles and the phenotype of flowering time from both the allele phasing workflow (first row) and marker analysis (second row) based on days after January 1^st^ on the median of the years 1999 and 2009-2016. Marker analysis was performed in 151 F_1_ individuals of the population GF.GA-47-42 x ‘Villard Blanc’, while allele phasing was performed in 35 F_1_ individuals. Number of alleles over the analyzed F_1_ individuals and the median of each, are given in the same order as in column 3. ab x cd: four alleles/both parents heterozygous, ef x eg: 3 alleles/both parents heterozygous, lm x ll: 2 alleles/ mother heterozygous, nn x np: 2 alleles, father heterozygous.

### Analysis of gene expression kinetics

Variation in expression could be detected in both time courses 2012/2013 and 2013/2014 for various FTC candidate and target genes when testing for time-specific effects. Between consecutive developmental stages of bud differentiation before dormancy (August 2^nd^ to September 5^th^, 2013 time series 1, Table 2) differences in expression could be detected for the MADS transcription factor *VvTM8* as well as the protein kinase encoding gene Vv*WNK5. VvTM8* encodes a MIKC transcription factor whose *A. thaliana* homologue *AtTM8* has been shown to be involved in the specification of flower organ identity [25].

In a time course of dormant buds (BBCH 0) until after bud burst when leaf formation had already begun (BBCH 11-13), 58 of the FTC candidate genes were found to show a BBCH or developmental stage-dependent expression. Several of these genes are squamosa binding proteins, MADS-and MYC transcription factors that are known to influence floral development. Most of these genes show a variation in gene expression due to an up or down regulation towards developmental stages during inflorescence maturation. In order to test for expression variation between consecutive developmental stages of bud development before inflorescence structures become externally visible, inflorescences collected after bud break were excluded from the analysis. Genes with different expression kinetics when the time course was extended to include visible inflorescences, are those showing a clear variation in gene expression between buds and inflorescence. In total 67 of such “inflorescence-specific genes” were identified (S8 Table).

After excluding inflorescences, several genes were found showing an obvious time-dependent expression. They cluster into two groups: genes upregulated in winter during bud dormancy (Fig 4, upper part) and genes upregulated towards inflorescence development (Fig 4, lower part). Most of these genes encode BZIP-, MADS-or MYC-transcription factors, which regulate other flowering related genes. Downregulation towards bud burst and inflorescence maturation was found for transcription factor genes involved in circadian rhythm such as *VvGRP2A (Glycine Rich Protein 2A), VvRVE1 (REVEILLE), VvTICb* (*TIME FOR COFFEE*) and *VvELF3* (*EARLY FLOWERING3*). Moreover, genes coding for transcription factors involved in gibberellic acid (GA) biosynthesis were found to be upregulated during bud dormancy. Numerous other genes like *VvHUA2b (ENHANCER OF AGAMOUS)*, which is involved in the repression of floral transition and flower development, were found to be upregulated during bud dormancy.

**Fig 4:**
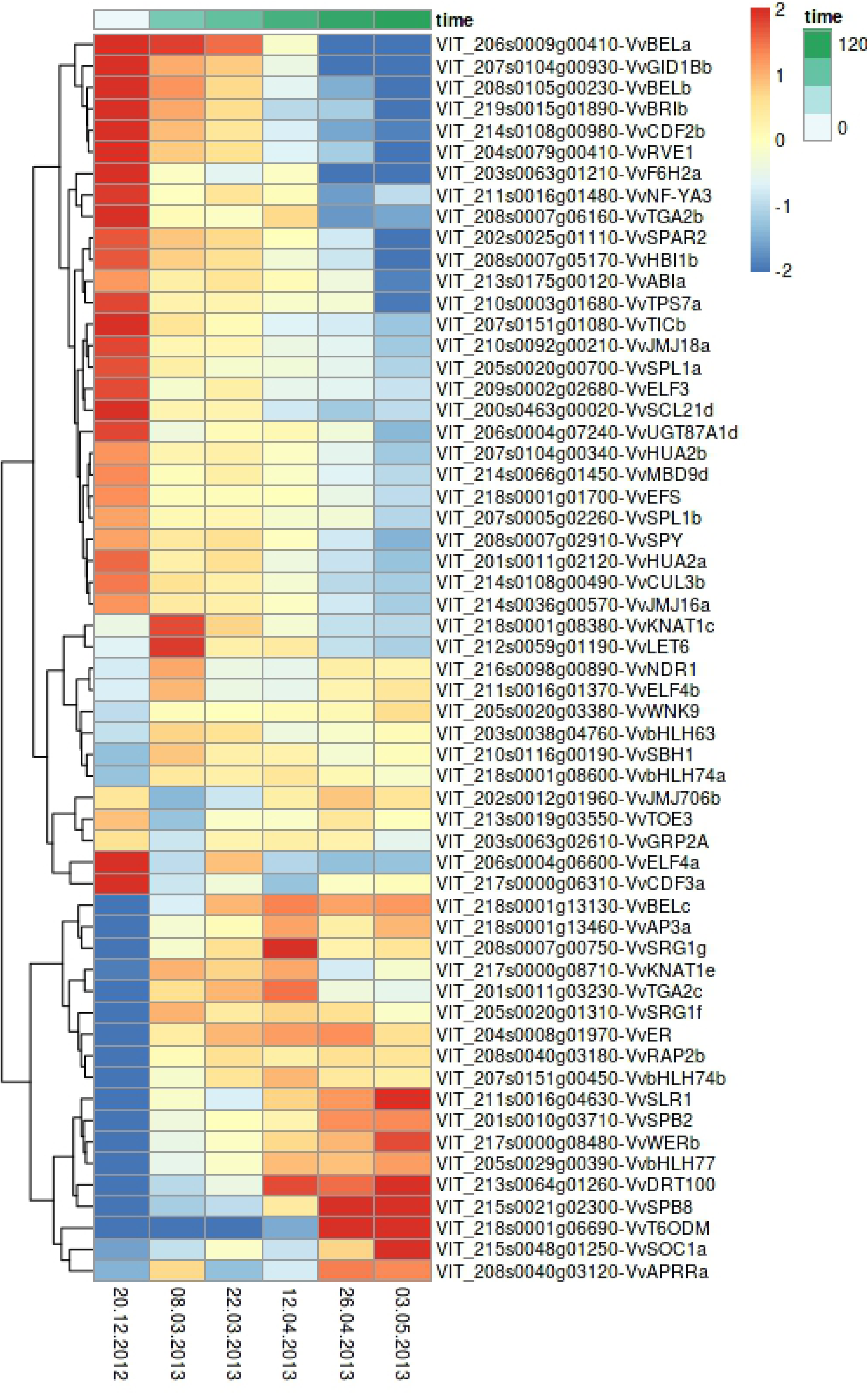
Heatmap of FTC candidate genes showing variations in their expression over consecutive time points of bud development from dormancy until appearance of inflorescence in grapevine variety GF.GA-47-42. Time series from December 20^th^, 2012 to May 3^rd^, 2013. LFC-threshold: 2 = expression fourfolded, −2 = expression quartered. Shown are rlog transformed counts.

For most of the genes (Fig 4) an up-or downregulation in expression is observed between the first and the second time point during bud dormancy. Many genes also show an up-or downregulation in expression between the third and the fourth time point when swelling buds are developing.

The gene expression for the amplicon sequenced target genes in buds and inflorescences is shown in Fig 5. Some genes are not expressed at all, while some are only expressed before dormancy or in inflorescence tissue. However, up-or downregulation in gene expression mainly occurs when swelling buds develop. Genes involved in floral development, such as *VvSEP3* and *4, VvAP1*, and *VvTM6* show an increased expression in developing inflorescences. *VvTM6* is a MADS-box B-class floral identity gene influencing the development of petals and stamen [52,53]. In *Vitis* all three B-class floral homeotic genes (*VvPI, VvAP3* and *VvTM6*) are highly expressed in inflorescences (S3 Figure).

**Fig 5:**
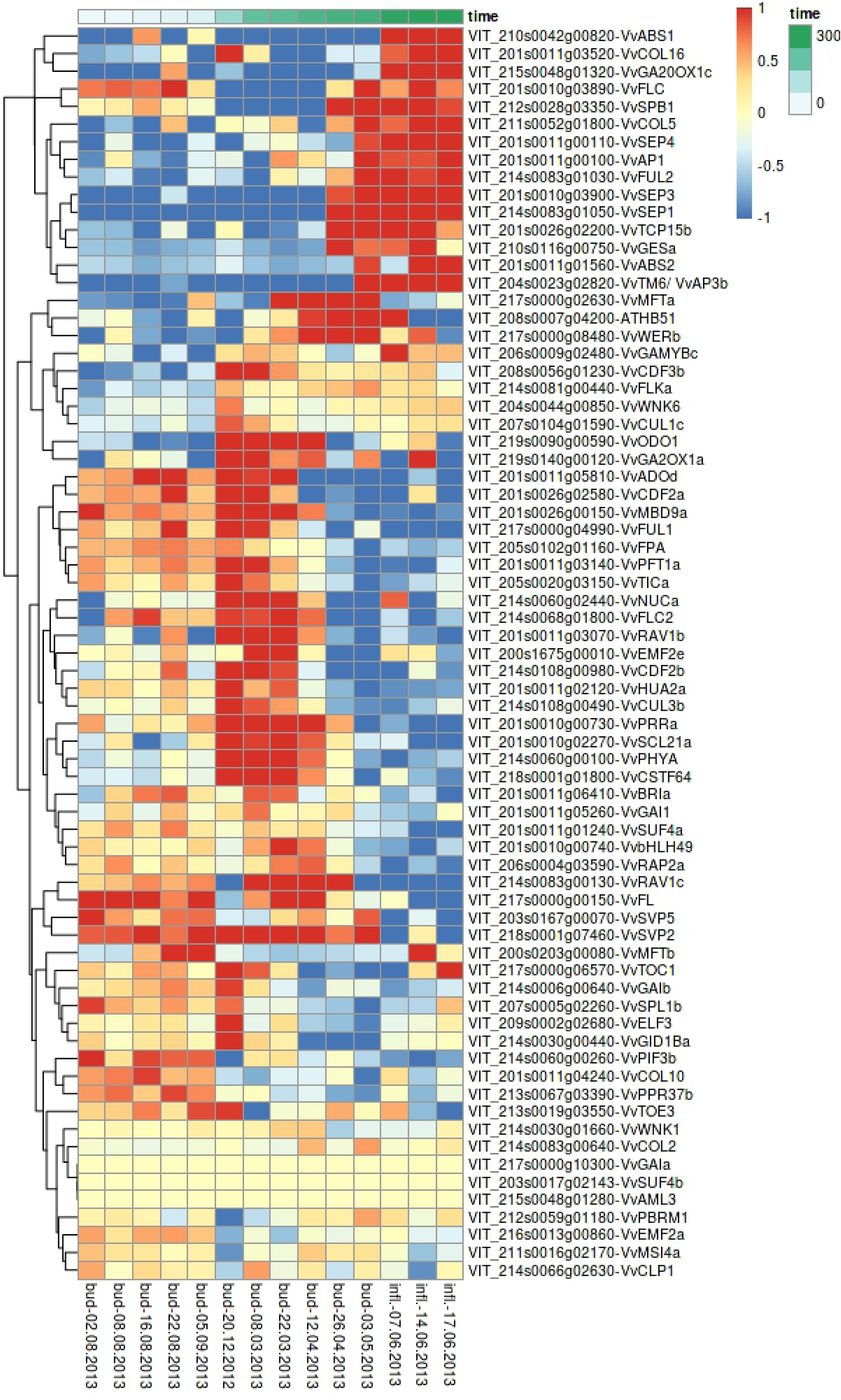
Heatmap of gene expression of amplicon sequenced FTC candidate genes in GF.GA-47-42 at different developmental stages of buds and inflorescences. LFC-threshold: 1 = expression doubled, −1 = expression halved. rlog transformed counts are shown.

For three selected time points, bud/inflorescence samples and the corresponding leaf from the same node were collected and differential gene expression was analyzed between leaves and the associated bud/inflorescence. Fig 6 shows a heatmap of the FTC candidate genes with expression differences between leaves and buds/inflorescences. With few exceptions, all genes with expression differences between leaves and buds or inflorescences are downregulated or not expressed in leaves.

**Fig 6:**
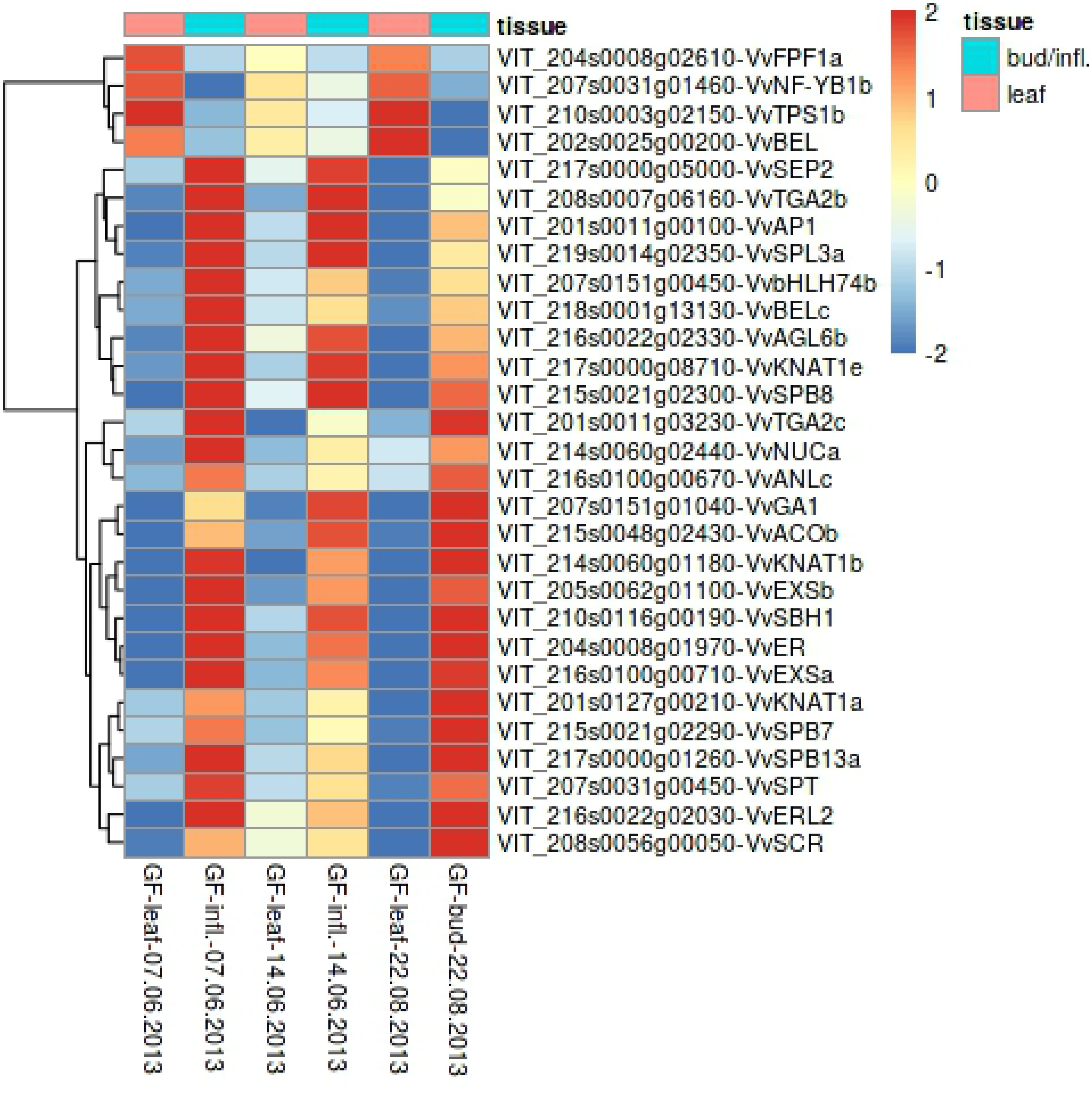
Heatmap of FTC candidate genes showing expression variations between leaves and their prompt buds/ inflorescences. LFC-threshold: 2 = expression fourfolded, −2 = expression quartered. Shown are rlog transformed counts.

## Discussion

### FTC candidate genes

A large number of FTC candidate genes inside and outside of known flowering QTLs in grapevine were identified. Although the identification relies mostly on sequence homology to previously known genes from other plants, the putative functional connection via e.g. Pfam, literature search or the performed RNA-Seq experiments substantiate the reliability of the prediction. This comprehensive gene list opens the door for investigations on e.g. flowering time networks in the future. One the one hand, compared to *Arabidopsis thaliana* there is probably an overestimation of FTC candidate genes in *Vitis*. On the other hand the high complexity and long duration of bud initiation and flower development may require a large number of genes.

### Allele phasing of target genes

A workflow for the phasing of amplicon sequenced genes using Illumina short-read sequencing of a diploid organism was established and successfully applied to separate alleles in regions with a length of up to 8.3 kb. By analyzing inheritance patterns within a family of parents and F_1_ individuals, we could show that the inheritance of alleles of neighboring genes within a QTL remains largely constant throughout the QTL. Since grapevine has a highly heterozygous genome and suffers from inbreeding depression, we used a F_1_ mapping population and followed a double pseudo-testcross strategy [54]. Therefore, a lower recombination frequency was expected compared to typical F_2_ mapping populations in other plant species. The constancy of the inheritance pattern of alleles of closely neighboring genes indicates the functionality and applicability of the established allele phasing method.

For the phasing of alleles, a mapped read or read pair needs to encompass two or more heterozygous sequence positions. The phase of the heterozygous sequence positions can be determined since each read or pair of reads is obtained from a single haplotype. Read lengths after trimming was distributed between 80 and 300 bp with an average insert size of ∼500 bp. When variants were located farther apart than the maximum length that could be spanned by a read pair, alleles could not be phased despite the presence of variants. Moreover, the allele frequency, calculated from the read coverage of variants can vary despite being amplified from the same allele. The amount of reads covering a variant can differ from one variant to the next. When dealing with extremely biased allele frequencies, this can lead to some variants being detected while others remain undetected. In such cases allele phasing was unsuccessful. Some amplicons could hardly be amplified at all. This is likely due to a high diversity at the primer binding sites between the reference sequence and the plant lines analyzed in this work.

The use of paired-end sequencing is highly advantageous in haplotype phasing as it covers variants that are spaced at distances longer than the technology’s read length limit. Read length in high-throughput sequencing is constantly increasing and technologies are evolving rapidly. With the rise of third generation technologies, capable of producing even longer reads, many of the difficulties associated with haplotype phasing might soon be alleviated as such data may permit direct phasing from sequence reads [26].

### Correlation analysis

We were able to detect a correlation between alleles of FTC target genes and flowering time for several QTL regions, which supports the role of these regions in the timing of flowering. Flowering time is highly dependent on the weather conditions of the respective and previous year. Therefore, correlation values vary between the years, as observed e.g., for Vv*WNK6* in 2016 (Table 3).

Alleles of FTC target genes within a QTL region on chr 1 were found to rather correlate with late, while QTL regions on chr 4 and 14 were found to correlate with early flowering. With one exception, all analyzed F_1_ individuals carrying alleles correlating with flowering time from two of the QTL regions on chr 1, 4, and 14 or all three of them were either intermediate-early, early, or very early flowering. The correlation for the QTL regions on chr 4 and 14 was more stable than for chr 1 indicating a stronger affect of these QTLs in the timing of flowering. The investigation of epistatic effects between these QTL regions could contribute to the clarification of the genetic factors that influence and control flowering time in grapevine.

Correlation values between alleles of FTC target genes and flowering time phenotypes could be largely supported by genetic marker analysis. Deviations can be due to the measuring method that can occasionally lead to deviations of up to two bp in product size. In order to distinguish the maximum putative number of alleles at a single locus within a bi-parental F_1_ population of a diploid organism, the marker needs to be capable of distinguishing between four different alleles.

Classic high informative marker analysis requires InDels / SSRs that distinguish between the maximum number of different alleles with polymorphic differences of at least two bp in size at a specific locus. The usage of blocks of tightly linked polymorphisms and treating each haplotype of these blocks as a separate allele can produce highly polymorphic markers. In addition, it also uses SNPs and InDels shorter than two bp to distinguish between the alleles. This leads to a higher resolution compared to classic marker analysis and the detection of a higher number of different alleles.

The correlation of alleles of FTC genes with flowering time phentoypes is based on the genotypic data on one hand, which is obtained through the allele phasing workflow, from amplicon sequencing, mapping and variant calling to the final establishment of allele sequences. On the other hand, the correlation analysis is based on the phenotypic data, which is also prone to errors. Phenotyping of flowering time was performed on a daily basis throughout the flowering phase. Differences in the timing of flowering shorter than one day are therefore not recorded. Moreover, phenotyping is a subjective process when different people work on the recording of phenotypic data and hence a possible error source.

As already mentioned, the timing of flowering depends clearly on environmental parameters, especially weather and climatic conditions. These are most probably non-genetic factors causing the differences in the flowering periods between the respective years. In 2016, for example, flowering in the population GF.GA-47-42 x ‘Villard Blanc’ started on June 17^th^ being very late compared to other years (Table 2). However, the flowering period was very short, ending after only 10 days on June 26^th^. Global radiation is distributed between ∼502 and ∼536 KWh/m^2^ at the beginning of flowering in the analyzed population and between ∼548 and ∼597 KWh/m^2^ at the end of it. While flowering occurred very late in 2016 compared to other years, the amount of global radiation until the first day of the flowering period was less than in the other years. This shows that the amount of solar radiation before flowering initiation was small which might have had an impact on the timing of flowering.

In some cases the p-value of correlation is significant although the medians are nearly equal or equal. This is because the Wilcoxon Rank-Sum test is a rank sum tests and not a median test. It ranks all of the observations from both groups and then sums the ranks from one of the groups and compares it with the expected rank sum. Therefore, it is in rare cases for groups possible to have different rank sums and yet have equal or nearly equal medians.

*VvHUA2a* of which an amplicon sequenced allele from ‘Villard Blanc’ was found to correlate with late flowering is a floral homeotic gene. It’s homologue in *A. thaliana, HUA2*, regulates the expression of the floral homeotic class-C gene *AGAMOUS (AG)* and *FLC [55]*. This suggests a role of *VvHUA2* in the delay of flowering.

An allele of *VvGAI1* from late flowering ‘Villard Blanc’ was found to correlate with late flowering. Mutants of *VvGAI1* are insensitive to gibberellic acid and form inflorescences instead of tendrils. These mutants show a correlation between inflorescence development and increased *VvFL* expression, a floral developmental gene [56]. In *A. thaliana, GAI* acts as a repressor of *LFY* and *SOC1* and thus represses flowering.

From the amplicon sequenced and early flowering individuals (median data) of the population GF.GA-47-42 x ‘Villard Blanc’, 90% were found to carry the *VvTM6* E1 allele inherited from GF.GA-47-42. Only 10% of plants that carry the other maternal allele are early flowering. *VvTM6* is a MADS-box B-class floral identity gene and influences the development of petals and stamen. In *A. thaliana*, mutants exhibit a transformation of petals to sepals and stamen to carpels. B-class floral homeotic genes either belong to the paleoAPETALA3 or to the PISTILLATA (PI) gene lineage, which are paralogous and resulted from a duplication event before the emergence of angiosperms. The paleoAP3 lineage underwent a further duplication event at the base of the core eudicots resulting in the two sublineages *euAP3* and *TM6* (named after the Tomato MADS-box gene 6) [57]. A *TM6* homologue is absent in *A. thaliana* [52,57,58]. In grapevine, all three B-class floral homeotic genes were found to be highly expressed in inflorescences (Fig 5) but not in leaves (Fig 6). [25] showed that *VvTM6 (VvAP3.2)* is expressed in fruits, while the expression of *VvAP3* (*VvAP3.1*) and *VvPI* is more restricted to flowers. Also, [52] showed that the expression of *VvTM6* is higher in carpels, fruits, and seeds than in petals. Due to the expression of *VvTM6* in carpels and during berry development and ripening, it was suggested to play an important role in grapevine fruit development [25]. The expression of *VvTM6* increases towards inflorescence maturation, which is followed by berry formation and ripening. This is consistent with its role during berry development and ripening.

All early flowering amplicon sequenced individuals of the population GF.GA-47-42 x ‘Villard Blanc’ were observed to carry the E1 allele of *VvWNK6* (Fig 3). In *A. thaliana WNK6* has been shown to be involved in circadian rhythm [59]. WNKs are a subfamily of serine/threonine protein kinases with a lysine residue essential for ATP-binding, which is located in kinase subdomain I instead of subdomain II as common among all other kinases [60]. It has been suggested that *WNK* gene family members regulate flowering time in *A. thaliana* by modulating the photoperiod pathway. For instance, APRR3, a component of the clock-associated APRR1/TOC1 quintet is a substrate of WNK1 in *A. thaliana*. T-DNA knockout mutants of *AtWNK1* are delayed in flowering time while T-DNA knockout mutants of *AtWNK2, 5*, and *8* flower early [61]. *WNK6* transcription is downregulated in *AtABI4* mutants, which show an early flowering phenotype [62]. In *A. thaliana, ABI4* negatively regulates flowering through directly promoting *FLC* transcription, a negative regulator of flowering [63]. This might indicate that *VvWNK6* is involved in the delay of flowering. *VvWNK6* expression was detected in leaves, buds, and inflorescences of the early flowering GF.GA-47-42. Both alleles E1 and E2 are expressed at a similar level. However, all individuals of the mapping population carrying the E1 allele of *VvWNK6* flower early. This suggests that either the E1 allele of *VvWNK6* itself might contribute to early flowering or alleles of other nearby-genes inherited together with E1 of *VvWNK6*. Further analysis should include the investigation of sequence variations leading to an alteration of the amino acid sequence and the functionality of the protein.

### Gene expression kinetics

Many of the analyzed FTC candidate genes show variations in expression pattern in the course of the developmental cycle, supporting their role in flowering time control. Genes coding for transcription factors and other proteins involved in inflorescence architecture, floral transition and flower development are usually upregulated after bud burst, while genes coding for proteins that repress flowering in diverse manners typically show an upregulation during bud dormancy (Fig 5). Among the genes showing downregulation towards bud burst and inflorescence maturation are transcription factors involved in circadian rhythm such as *VvGRP2A (Glycine Rich Protein 2A), VvRVE1* (*REVEILLE1*), *VvTICb* (*TIME FOR COFFEE*) and *VvELF3* (*EARLY FLOWERING3*). It is not unexpected to detect different gene expression kinetics for genes involved in circadian rhythm since sampling was performed at the same time of the day over the entire time course. However, the period from daybreak until the time of sampling varies throughout the year and the different seasons.

*AtGRP7*, the homologue of *VvGRP2A* in *A. thaliana*, undergoes circadian oscillations with peak levels in the evening [64]. *RVE* is a MYB-like transcription factor that controls auxin levels, promotes free auxin and hence plant growth during the day [65]. *TIC* and *ELF3* are components of the circadian clock in *A. thaliana*. *ELF3* is a circadian clock gene that contributes to photoperiod-dependent flowering in plants [66-68]. Our findings thus suggest a similar impact of these genes in grapevine.

Moreover, genes coding for transcription factors involved in GA biosynthesis were found to be upregulated during bud dormancy. GAs are inhibitors of flowering in many fruit species but their role in grapevine varies with the stage of bud development. The initiation and development of lateral meristems is promoted by GAs as well as their development into tendrils, while inflorescence development is suppressed by GAs. Thus GA is a promoter of flowering at an early stage but acts as an inhibitor of flowering later on and promotes vegetative growth [19]. *SPY (SPINDLY)*, whose *Vitis* homologue *VvSPY* was found to be upregulated during bud dormancy, is a negative regulator of GA response in *A. thaliana* and functions with *GI (GIGANTEA)* in pathways controlling flowering [69]. In *Vitis* the role of *SPY* in GA signaling is still unclear. It could be shown that treatment of grapevine plants at pre-bloom stage with GA led to rachis elongation and a downregulation of *VvSPY* in the rachis [70]. In *A. thaliana* GA signaling is initiated through its binding to the GA INSENSITIVE DWARF1 (GID1) receptors. This allows subsequent interaction between GID1 and DELLA proteins (GA INSENSITIVE [GAI], REPRESSOR OF GAI-3 [RGA], RGA-LIKE1 [RGL1], RGL2, and RGL3). DELLA proteins are transcriptional repressors and downregulate GA response genes. In the presence of gibberellin, the stable GID1-GA-DELLA complex is recognized by the SCF^SLY1^ complex which ubiquintylates the DELLA proteins and causes their degradation by the 26S proteasome [71,72]. It has been reported previously that GID1-transcripts are upregulated during bud dormancy in grapevine while transcripts of DELLA are downregulated [73]. Similarly, we found that the GID1B receptor transcript is upregulated during bud dormancy while the DELLA-protein SLR1-like (SLENDER RICE 1 LIKE) are downregulated. This confirms the promoting role of GID1B in plant growth, and the development of lateral meristems in dormant buds and indicates that SLR1-like is responsive for the mediation of the suppression of inflorescence development through GA.

In our analyses, numerous other genes involved in the repression of floral transition and flower development were found to be upregulated during bud dormancy. *HUA2*-like genes, which play a role in the repression of floral transition [74], are upregulated during bud dormancy in *Vitis*. The *KNOTTED1*-like homeobox gene *BP (BREVIPEDICELLUS)* was found to be upregulated towards grapevine bud burst and inflorescence maturation. In *A. thaliana BP* controls distal pedicel growth and thus inflorescence architecture [75,76]. *ER (ERECTA)* and other *KNAT (KNOTTED-LIKE)* genes, are involved in inflorescence architecture in *A. thaliana* [77,78], were also found to be upregulated towards bud burst, which indicates their function in inflorescence development. Genes for SQUAMOSA promoter-binding proteins, known to be involved in flower development [79], were downregulated during bud dormancy while upregulated during flower formation in grapevine. The *BEL*-like gene (*VvBELa* and *b*) and the *Vitis STM* orthologue *VvSBH1* were also found to be upregulated during bud dormancy. STM and the *A. thaliana* homeobox-gene *BEL1* build a complex, which maintains the indeterminacy of the inflorescence meristem [80].

MYC transcription factors *VvbHLH74* and *VvbHLH63* show large variations in gene expression over time with a peak in expression around March when buds are swelling. *CIB1 (cryptochrome-interacting basic-helix-loop-helix)*, the *A. thaliana* homologue of *VvbHLH63*, plays a role in CRY2 (cryptochrome 2)-dependent regulation of flowering time. Cryptochromes (CRY) are blue-light receptors that mediate light response. In yeast and *A. thaliana*, CIB1 interacts with CRY2 when blue light is available. It promotes CRY2-dependent floral initiation together with additional CIB1-related proteins and stimulates *FT* transcription [81]. Hence, *VvbHLH74* and *VvbHLH63* might be involved in light dependent floral initiation.

*ELF*-like genes as well as a *CONSTANS*-like gene (*VvCOL16*) and *CDF* genes (*CYCLING DOF FACTORS*) were upregulated during bud dormancy. DOF proteins delay flowering by repressing *CO* transcription [82]. *ELF3, ELF4*, and *TOC1* function in the primary, phytochrome-mediated light-input pathway to the circadian oscillator in *A. thaliana. TOC1* is necessary for light-induced *CCA1* (CIRCADIAN CLOCK ASSOCIATED 1)/ LHY (*LATE ELONGATED HYPOCOTYL*) expression [83]. Mutants of *elf4* show attenuated expression of *CCA1* and early flowering in non-inductive photoperiods, which is probably caused by elevated amounts of *CONSTANS (CO)*, a gene that promotes floral induction [84]. *ELF4* is a flowering pathway gene that may play a key role in signaling processes regulating dormancy induction in grapevine [85].

*MBD9*, whose *A. thaliana* homologue -*AtMBD9* is related to the inhibition of flowering [86] and suggested to have a role in bud development through an interaction with FLC in *A. thaliana* [85], is upregulated during bud dormancy in grapevine. *VvSPAR2* (*SUPPRESSOR OF PHYA RELATED2*) is upregulated during bud dormancy and downregulated towards inflorescence development. Its homologue in *A. thaliana* represses photomorphogenesis by negatively regulating the transcription factor *HY5 (ELONGATED HYPOCOTYL 5)*, which promotes photomorphogenesis [87,88].

## Conclusion

Here, we have reported a new workflow for amplicon sequencing including allele phasing in the highly heterozygous species grapevine. Our genetic association study revealed a significant correlation between alleles of selected FTC target genes and flowering time phenotypes within and outside of previously mapped QTL regions for flowering time on chr 1, 4, 14, 17, and 18. The discovery of a correlation between alleles of FTC target genes and the timing of flowering for genes within previously defined QTL regions supports the role of these QTLs in the timing of flowering. The analysis of gene expression kinetics revealed strong changes in expression pattern for many FTC candidate genes over the consecutive developmental stages. A shift between an up-or downregulation in expression mostly occurred between dormant and swelling buds, or toward inflorescence maturation when the young inflorescence structures at the shoots grow out of the buds and become externally visible. These time-dependent expression profiles underline the role of many FTC candidate genes in the control of flowering time. Moreover, many FTC candidate genes were found to be expressed in buds and inflorescences but not in leaves. This tissue specificity further confirms their role in flowering time and floral development.

The knowledge of genes and loci that influence flowering time and play a role in early flowering may allow the selection of genotypes not carrying these alleles through grapevine breeding programs. To meet the expected change of climate conditions late flowering cultivars might be better adapted, especially in the present cool climate areas.

For future research, grapevine cultivars are to be analysed for alleles of flowering time control genes correlating with early or late flowering in order to further investigate the role of these alleles in the timing of flowering and study epistatic and additive effects between QTL regions influencing the timing of flowering.

## Author contributions

N.K., L.H. and D.H. concieved and planned the experiments. N.K., I.O., A.S., P.V. designed and performed the experiments. I.O., L.H. and A.S. performed the phenotyping. N.K., I.O. and P.V. carried out the sample preparation. N.K., I.O. and L.H. calculated the data. R.T., B.W., D.H. concieved the original idea and supervised the project. All authors interpreted and discussed results. N.K. and D.H. wrote the manuscript with input from all authors.

## Acknowledgements

We thank Willy Keller, Johanna Müschner, Thomas Rosleff Sörensen, and Andreas Preiss for the excellent technical support, and Martin Sagasser for editing the manuscript. In addition, the authors wish to thank the members of the SPP1530 team for their support. The research was funded by the German research foundation (DFG) - Project number 172418080/15-SPP1530.

## Bibliography

## Supplementary Figures

S1 Figure: Illustration of the reproductive developmental cycle of grapevine showing the stages of flowering and berry development ([according to 1]). UP: uncommitted primordia.

S2 Figure: Flowering periods in days after January 1^st^ in the population GF.GA-47-42 x ‘Villard Blanc’ in the years 1999 and 2010 – 2016.

S3 Figure: Expression profile of the three B-class floral homeotic genes *VvAP3, VvTM6* and *VvPI* over consecutive developmental stages of bud-and inflorescence development in GF.GA-47-42. The last three time points refer to developing stages of visible inflorescence structures.

S1 Table: F_1_ individuals of the mapping population Gf.Ga-47-42 x Villard blanc and days until full bloom after January 1st in the years 1999, 2009, and 2011-2016.

S2 Table: Median flowering time of the years 1999 and 2010-2016 for amplicon sequenced F_1_ individuals.

S3 Table: Nomenclature system for the alleles of genes. S4 Table: FTC candidate genes in Vitis.

S5 Table: Amplicon sequenced FTC target genes; genomic positions and length of phased intervals. S6 Table: Molecular marker information.

S7 Table: Correlation analysis with molecular markers for F1 mapping population. S8 Table: Inflorescence-specific FTC candidate genes.

## References

1. Carmona MJ, Chaïb J, Martínez-Zapater JM, Thomas MR (2008) A molecular genetic perspective of reproductive development in grapevine. Journal of Experimental Botany 59: 2579–2596.

2. Duchêne E, Butterlin G, Dumas V, Merdinoglu D (2012) Towards the adaptation of grapevine varieties to climate change: QTLs and candidate genes for developmental stages. Theoretical and Applied Genetics 124: 623–635.

3. Mullins MG, Bouquet A, Williams LE (1992) Biology of the Grapevine. Cambridge: Cambridge University Press.

4. Tomasi D, Jones GV, Giust M, Lovat L, Gaiotti F (2011) Grapevine Phenology and Climate Change: Relationships and Trends in the Veneto Region of Italy for 1964-2009. American Journal of Enology and Viticulture 62: 329–339.

5. Zyprian A, Eibach R, Trapp O, Schwander F, Töpfer R (2018) Grapevine breeding under climate change: Applicability of a molecular marker linked to veraison. Vitis 57: 119–123.

6. Rienth M, Torregrosa L, Sarah G, Ardisson M, Brillouet JM, et al. (2016) Temperature desynchronizes sugar and organic acid metabolism in ripening grapevine fruits and remodels their transcriptome. BMC Plant Biology 16: 164.

7. Boss PK, Buckeridge EJ, Poole A, Thomas MR (2003) New insights into grapevine flowering. Functional Plant Biology 30: 593–606.

8. Andrés F, Coupland G (2012) The genetic basis of flowering responses to seasonal cues. Nature Reviews Genetics 13: 627–639.

9. Ehrenreich IM, Hanzawa Y, Chou L, Roe JL, Kover PX, et al. (2009) Candidate gene association mapping of Arabidopsis flowering time. Genetics 183: 325–335.

10. Jaillon O, Aury JM, Noel B, Policriti A, Clepet C, et al. (2007) The grapevine genome sequence suggests ancestral hexaploidization in major angiosperm phyla. Nature 449: 463–467.

11. Velasco R, Zharkikh A, Troggio M, Cartwright DA, Cestaro A, et al. (2007) A high quality draft consensus sequence of the genome of a heterozygous grapevine variety. PLoS One 2.

12. Vitulo N, Forcato C, Carpinelli EC, Telatin A, Campagna D, et al. (2014) A deep survey of alternative splicing in grape reveals changes in the splicing machinery related to tissue, stress condition and genotype. BMC Plant Biology 14: 99.

13. Adam-Blondon A-F, Jaillon O, Vezzulli S, Zharkikh A, Troggio M, et al. (2011) Genome Sequence Initiatives. In: Adam-Blondon A-F, Martinez-Zapater JM, Kole C, editors. Genetics, Genomics and Breeding of Grapes.

14. Canaguier A, Grimplet J, Di Gaspero G, Scalabrin S, Duchêne E, et al. (2017) A new version of the grapevine reference genome assembly (12X.v2) and of its annotation (VCost.v3). Genomics Data 14: 56–62.

15. Carmo Vasconcelos M, Greven M, Winefield CS, Trought MCT, Raw V (2009) The Flowering Process of Vitis vinifera: A Review. American Journal of Enology and Viticulture 60: 411–434.

16. Calonje M, Cubas P, Martínez-Zapater JM, Carmona MJ (2004) Floral meristem identity genes are expressed during tendril development in grapevine. Plant Physiology 135: 1491–1501.

17. Joly D, Perrin M, Gertz C, Kronenberger J, Demangeat G, et al. (2004) Expression analysis of flowering genes from seedling-stage to vineyard life of grapevine cv. Riesling. Plant Science 166: 1427–1436.

18. Sreekantan L, Thomas MR (2006) VvFT and VvMADS8, the grapevine homologues of the floral integrators FT and SOC1, have unique expression patterns in grapevine and hasten flowering in Arabidopsis. Functional Plant Biology 33: 1129–1139.

19. Carmona MJ, Cubas P, Calonje M, Martinez-Zapater JM (2007) Flowering transition in grapevine (Vitis vinifera L.). Canadian Journal of Botany 85: 701–711.

20. Grattapaglia D, Sederoff R (1994) Genetic linkage maps of Eucalyptus grandis and Eucalyptus urophylla using a pseudo-testcross: mapping strategy and RAPD markers. Genetics 137: 1121–1137.

21. Cipriani G, Di Gaspero G, Canaguier A, Jusseaume J, Tassin J, et al. (2011) Molecular Linkage Maps: Strategies, Resources and Achievements. In: Adam-Blondon A-F, Martinez-Zapater JM Kole C, editors. Genetics, Genomics and Breeding of Grapes.

22. Costantini L, Battilana J, Lamaj F, Fanizza G, Grando MS (2008) Berry and phenology-related traits in grapevine (Vitis vinifera L.): from quantitative trait loci to underlying genes. BMC Plant Biology 8.

23. Fechter I, Hausmann L, Zyprian E, Daum M, Holtgräwe D, et al. (2014) QTL analysis of flowering time and ripening traits suggests an impact of a genomic region on linkage group 1 in Vitis. Theoretical and Applied Genetics 127: 1857–1872.

24. Gökbayrak Z, Özer C, Söylemezoglu G (2006) Preliminary Results on Genome Mapping of an Italia x Mercan Grapevine Population. Turk J Agric For 30: 273–280.

25. Díaz-Riquelme J, Lijavetzky D, Martínez-Zapater JM, Carmona MJ (2009) Genome-wide analysis of MIKCC-type MADS box genes in grapevine. Plant Physiology 149: 354–369.

26. Aguiar D, Istrail S (2012) HapCompass: a fast cycle basis algorithm for accurate haplotype assembly of sequence data. Journal of Computational Biology 19: 577–590.

27. Browning SR, Browning BL (2011) Haplotype phasing: existing methods and new developments. Nature Reviews Genetics 12: 703–714.

28. Keurentjes JJ, Fu J, De Vos RCH, Lommen A, Hall RD, et al. (2006) The genetics of plant metabolism. Nature Genetics 38: 842–849.

29. Martin M, Patterson M, Garg S, Fischer SO, Pisanti N, et al. (2016) WhatsHap: fast and accurate read-based phasing. Software.

30. Bansal V, Bafna V (2008) HapCUT: an efficient and accurate algorithm for the haplotype assembly problem. Bioinformatics 24: 153–159.

31. Lancia G, Bafna V, Istrail S, Lippert R, Schwartz R (2001) SNPs Problems, Complexity, and Algorithms. ESA: 182–193.

32. Zyprian E, Ochßner I, Schwander F, Šimon S, Hausmann L, et al. (2016) Quantitative trait loci affecting pathogen resistance and ripening of grapevines. Molecular Genetics and Genomics 291: 1573–1594.

33. Lorenz DH, Eichhorn KW, Bleiholder H, Klose R, Meier U, et al. (1995) Phenological growth stages of the grapevine (Vitis vinifera L. ssp. vinifera)-Codes and descrptions according to the extended BBCH scale. Australian Journal of Grape and Wine Research 1: 100–110.

34. Camacho C, Coulouris G, Avagyan V, Ma N, Papadopoulos J, et al. (2009) BLAST+: architecture and applications. BMC Bioinformatics 10: 421.

35. Ward N, Moreno-Hagelsieb G (2014) Quickly finding orthologs as reciprocal best hits with BLAT, LAST, and UBLAST: how much do we miss? PLoS ONE 9: e101850.

36. Untergasser A, Cutcutache I, Koressaar T, Ye J, Faircloth BC, et al. (2012) Primer3--new capabilities and interfaces. Nucleic Acids Research 40.

37. Bolger AM, Lohse M, Usadel B (2014) Trimmomatic: a flexible trimmer for Illumina sequence data. Bioinformatics 30: 2114–2120.

38. Li H, Durbin R (2009) Fast and accurate short read alignment with Burrows-Wheeler transform. Bioinformatics 25: 1754–1760.

39. Li H, Handsaker B, Wysoker A, Fennell T, Ruan J, et al. (2009) The Sequence Alignment/Map format and SAMtools. Bioinformatics 25: 2078–2079.

40. McKenna A, Hanna M, Banks E, Sivachenko A, Cibulskis K, et al. (2010) The Genome Analysis Toolkit: a MapReduce framework for analyzing next-generation DNA sequencing data. Genome Research 20: 1297–1303.

41. Van der Auwera GA, Carneiro MO, Hartl C, Poplin R, Del Angel G, et al. (2013) From FastQ data to high confidence variant calls: the Genome Analysis Toolkit best practices pipeline. Current Protocols in Bioinformatics 11: 1110.

42. DePristo MA, Banks E, Poplin R, Garimella KV, Maguire JR, et al. (2011) A framework for variation discovery and genotyping using next-generation DNA sequencing data. Nature Genetics 43: 491–498.

43. Kim D, Pertea G, Trapnell C, Pimentel H, Kelley R, et al. (2013) TopHat2: accurate alignment of transcriptomes in the presence of insertions, deletions and gene fusions. Genome Biology 14: R36.

44. Anders S, Pyl PT, Huber W (2015) HTSeq--a Python framework to work with high-throughput sequencing data. Bioinformatics 31: 166–169.

45. Love MI, Huber W, Anders S (2014) Moderated estimation of fold change and dispersion for RNA-seq data with DESeq2. Genome Biology 15: 550.

46. Riechmann JL, Meyerowitz EM (1998) The AP2/EREBP family of plant transcription factors. Biological Chemistry 379: 633–646.

47. Gehring WJ (1992) The homeobox in perspective. Trends in Biochemical Sciences 17: 277–280.

48. Stracke R, Werber M, Weisshaar B (2001) The R2R3-MYB gene family in Arabidopsis thaliana. Current Opinion in Plant Biology 4: 447–456.

49. Johanson U, West J, Lister C, Michaels S, Amasino R, et al. (2000) Molecular analysis of FRIGIDA, a major determinant of natural variation in Arabidopsis flowering time. Science 290: 344–347.

50. Pysh LD, Wysocka-Diller JW, Camilleri C, Bouchez D, Benfey PN (1999) The GRAS gene family in Arabidopsis: sequence characterization and basic expression analysis of the SCARECROW-LIKE genes. The Plant Journal 18: 111–119.

51. Chen X, Zhang Z, Liu D, Zhang K, Li A, et al. (2010) SQUAMOSA promoter-binding protein-like transcription factors: star players for plant growth and development. Journal of Integrative Plant Biology 52: 946–951.

52. Poupin MJ, Federici F, Medina C, Matus JT, Timmermann T, et al. (2007) Isolation of the three grape sub-lineages of B-class MADS-box TM6, PISTILLATA and APETALA3 genes which are differentially expressed during flower and fruit development. Gene 404: 10–24.

53. Theissen G, Becker A, Di Rosa A, Kanno A, Kim JT, et al. (2000) A short history of MADS-box genes in plants. Plant Molecular Biology 42: 115–149.

54. Grattapaglia D, Bertolucci FL, Sederoff RR (1995) Genetic mapping of QTLs controlling vegetative propagation in Eucalyptus grandis and E. urophylla using a pseudo-testcross strategy and RAPD markers. Theoretical and Applied Genetics 90: 933–947.

55. Chen X, Meyerowitz, E M (1999) HUA1 and HUA2 are two members of the floral homeotic AGAMOUS pathway.. Molecular Cell 3: 349–360.

56. Boss PK, Thomas MR (2002) Association of dwarfism and floral induction with a grape ‘green revolution’ mutation. Nature 416: 847–850.

57. Causier B, Castillo R, Xue Y, Schwarz-Sommer Z, Davies B (2010) Tracing the evolution of the floral homeotic B-and C-function genes through genome synteny. Molecular Biology and Evolution 27: 2651–2664.

58. Kramer EM, Dorit RL, Irish VF (1998) Molecular evolution of genes controlling petal and stamen development: duplication and divergence within the APETALA3 and PISTILLATA MADS-box gene lineages. Genetics 149: 765–783.

59. Nakamichi N, Murakami-Kojima M, Sato E, Kishi Y, Yamashino T, et al. (2002) Compilation and characterization of a novel WNK family of protein kinases in Arabiodpsis thaliana with reference to circadian rhythms. Bioscience, Biotechnology, and Biochemistry 66: 2429–2436.

60. Hong-Hermesdorf A, Brüx A, Grüber A, Grüber G, Schumacher K (2006) A WNK kinase binds and phosphorylates V-ATPase subunit C. FEBS Letters 580: 932–939.

61. Wang Y, Liu K, Liao H, Zhuang C, Ma H, et al. (2008) The plant WNK gene family and regulation of flowering time in Arabidopsis. Plant Biology 10: 548–562.

62. Foyer CH, Kerchev PI, Hancock RD (2012) The ABA-INSENSITIVE-4 (ABI4) transcription factor links redox, hormone and sugar signaling pathways. Plant Signaling & Behavior 7: 276–281.

63. Shu K, Chen Q, Wu Y, Liu R, Zhang H, et al. (2016) ABSCISIC ACID-INSENSITIVE 4 negatively regulates flowering through directly promoting Arabidopsis FLOWERING LOCUS C transcription. Journal of Experimental Botany 67: 195–205.

64. Heintzen C, Nater M, Apel K, Staiger D (1997) AtGRP7, a nuclear RNA-binding protein as a component of a circadian-regulated negative feedback loop in Arabidopsis thaliana. Proceedings of the National Academy of Sciences of the United States of America 94: 8515–8520.

65. Rawat R, Schwartz J, Jones MA, Sairanen I, Cheng Y, et al. (2009) REVEILLE1, a Myb-like transcription factor, integrates the circadian clock and auxin pathways. Proceedings of the National Academy of Sciences of the United States of America 106: 16883–16888.

66. Boden SA, Weiss D, Ross JJ, Davies NW, Trevaskis B, et al. (2014) EARLY FLOWERING3 Regulates Flowering in Spring Barley by Mediating Gibberellin Production and FLOWERING LOCUS T Expression. The Plant Cell 26: 1557–1569.

67. Hall A, Bastow RM, Davis SJ, Hanano S, McWatters HG, et al. (2003) The TIME FOR COFFEE gene maintains the amplitude and timing of Arabidopsis circadian clocks. The Plant Cell 15: 2719–2729.

68. Ding Z, Millar AJ, Davis AM, Davis SJ (2007) TIME FOR COFFEE encodes a nuclear regulator in the Arabidopsis thaliana circadian clock. The Plant Cell 19: 1522–1536.

69. Tseng TS, Salomé PA, McClung CR, Olszewski NE (2004) SPINDLY and GIGANTEA interact and act in Arabidopsis thaliana pathways involved in light responses, flowering, and rhythms in cotyledon movements. The Plant Cell 16: 1550–1563.

70. Upadhyay A, Jogaiah S, Maske SR, Kadoo NY, Gupta VS (2015) Expression of stable reference genes and SPINDLY gene in response to gibberellic acid application at different stages of grapevine development. Biologia Plantarum 59: 436–444.

71. Cheng C, Jiao C, Singer SD, Gao M, Xu X, et al. (2015) Gibberellin-induced changes in the transcriptome of grapevine (Vitis labrusca × V. vinifera) cv. Kyoho flowers. BMC Genomics 16.

72. Fleet CM, Sun TP (2005) A DELLAcate balance: the role of gibberellin in plant morphogenesis. Current Opinion in Plant Biology 8: 77–85.

73. Khalil-Ur-Rehman M, Sun L, Li CX, Faheem M, Wang W, et al. (2017) Comparative RNA-seq based transcriptomic analysis of bud dormancy in grape. BMC Plant Biology 17: 18.

74. Doyle MR, Bizzell CM, Keller MR, Michaels SD, Song J, et al. (2005) HUA2 is required for the expression of floral repressors in Arabidopsis thaliana. The Plant Journal 41: 376–385.

75. Douglas SJ, Riggs CD (2005) Pedicel development in Arabidopsis thaliana: contribution of vascular positioning and the role of the BREVIPEDICELLUS and ERECTA genes. Developmental Biology 284: 451–463.

76. Venglat SP, Dumonceaux T, Rozwadowski K, Parnell L, Babic V, et al. (2002) The homeobox gene BREVIPEDICELLUS is a key regulator of inflorescence architecture in Arabidopsis. Proceedings of the National Academy of Sciences of the United States of America 99: 4730–4735.

77. Douglas SJ, Chuck G, Dengler RE, Pelecanda L, Riggs CD (2002) KNAT1 and ERECTA Regulate Inflorescence Architecture in Arabidopsis. The Plant Cell 14: 547–558.

78. Shpak ED, Berthiaume CT, Hill EJ, Torii KU (2004) Synergistic interaction of three ERECTA-family receptor-like kinases controls Arabidopsis organ growth and flower development by promoting cell proliferation. Development 131: 1491–1501.

79. Yamasaki K, Kigawa T, Inoue M, Yamasaki T, Yabuki T, et al. (2006) An Arabidopsis SBP-domain fragment with a disrupted C-terminal zinc-binding site retains its tertiary structure. FEBS Letters 580: 2109–2116.

80. Bellaoui M, Pidkowich MS, Samach A, Kushalappa K, Kohalmi SE, et al. (2001) The Arabidopsis BELL1 and KNOX TALE homeodomain proteins interact through a domain conserved between plants and animals. The Plant Cell 13: 2455–2470.

81. Liu H, Yu X, Li K, Klejnot J, Yang H, et al. (2008) Photoexcited CRY2 interacts with CIB1 to regulate transcription and floral initiation in Arabidopsis. Science 322: 1535–1539.

82. Fornara F, Panigrahi KC, Gissot L, Sauerbrunn N, Rühl M, et al. (2009) Arabidopsis DOF transcription factors act redundantly to reduce CONSTANS expression and are essential for a photoperiodic flowering response. Developmental Cell 17: 75–86.

83. Doyle MR, Davis SJ, Bastow RM, McWatters HG, Kozma-Bognár L, et al. (2002) The ELF4 gene controls circadian rhythms and flowering time in Arabidopsis thaliana. Nature 419: 74–77.

84. Kikis EA, Khanna R, Quail PH (2005) ELF4 is a phytochrome-regulated component of a negative-feedback loop involving the central oscillator components CCA1 and LHY. The Plant Journal 44: 300–313.

85. Sreekantan L, Mathiason K, Grimplet J, Schlauch K, Dickerson JA, et al. (2010) Differential floral development and gene expression in grapevines during long and short photoperiods suggests a role for floral genes in dormancy transitioning. Plant Molecular Biology 73: 191–205.

86. Peng M, Cui Y, Bi YM, Rothstein SJ (2006) AtMBD9: a protein with a methyl-CpG-binding domain regulates flowering time and shoot branching in Arabidopsis. The Plant Journal 46: 282–296.

87. Huang X, Ouyang X, Yang P, Lau OS, Chen L, et al. (2013) Conversion from CUL4-based COP1-SPA E3 apparatus to UVR8-COP1-SPA complexes underlies a distinct biochemical function of COP1 under UV-B. Proceedings of the National Academy of Sciences of the United States of America 110: 16669–16674.

88. Srivastava AK, Senapati D, Srivastava A, Chakraborty M, Gangappa SN, et al. (2015) Short Hypocotyl in White Light1 Interacts with Elongated Hypocotyl5 (HY5) and Constitutive Photomorphogenic1 (COP1) and Promotes COP1-Mediated Degradation of HY5 during Arabidopsis Seedling Development. Plant Physiology 169: 2922–2934.

